# Genealogical properties of subsamples in highly fecund populations

**DOI:** 10.1101/164418

**Authors:** Bjarki Eldon, Fabian Freund

**Affiliations:** Museum für Naturkunde 43 Invalidenstraße 10115 Berlin Germany; University of Hohenheim Institute 350b Fruwirthstraße 21 D-70599 Stuttgart, Germany

**Keywords:** coalescent, high fecundity, nested samples, multiple mergers, time to most recent common ancestor

## Abstract

We consider some genealogical properties of nested samples. The complete sample is assumed to have been drawn from a natural population characterised by high fecundity and sweepstakes reproduction (abbreviated HFSR). The random gene genealogies of the samples are — due to our assumption of HFSR — modelled by coalescent processes which admit multiple mergers of ancestral lineages looking back in time. Among the genealogical properties we consider are the probability that the most recent common ancestor is shared between the complete sample and the subsample nested within the complete sample; we also compare the lengths of ‘internal’ branches of nested genealogies between different coalescent processes. The results indicate how ‘informative’ a subsample is about the properties of the larger complete sample, how much information is gained by increasing the sample size, and how the ‘informativeness’ of the subsample varies between different coalescent processes.

## 1 Introduction

The study of the evolutionary history of natural populations usually proceeds by drawing inference from a random sample of DNA sequences. To this end the coalescent approach initiated by [56,58,57,83,52] - i.e. the probabilistic modeling of the random ancestral relations of the sampled DNA sequences - has proved to be very useful [85, cf.]. Inference based on the coalescent relies on the key assumption, as in standard statistical inference, that the evolutionary history of the (finite) sample approximates, or is informative about, the evolutionary history of the population from which the sample is drawn. We would like to know how much some basic genealogical sample-based statistics tell us about the population in a multiple-merger coalescent framework. Does the ‘informativeness’ of the various genealogical statistics depend on the underlying coalescent process? A more practical approach to this question is, instead of comparing a sample with the population, to ask how much of the genetic information of a sample is already contained in a subsample, i.e. what is gained by enlarging the sample? A related question concerns the size of the sample; i.e. how large does our sample need to be for a reliable inference? Do standard genetic approaches or guidelines, e.g. about sample size for population genetic studies, still hold true for populations characterised by high fecundity and sweepstakes reproduction (abbreviated HFSR)? In Sec. A1 we give a brief overview of population models of reproduction which admit HFSR. The coalescent processes derived from HFSR population models admit multiple-mergers of ancestral lineages (see Sec. A2).

We approach these problems by studying some genealogical properties of nested samples, by which we mean where a sample (a <u>subsample</u>) is drawn (uniformly at random without replacement) from a larger sample (the <u>complete</u> sample). By way of an example, [74] consider nested samples whose ancestries are governed by the Kingman *n*-coalescent [56,58,57]. One of the results of [74] concerns the probability that a subsample shares its <u>most recent common ancestor</u> (abbreviated MRCA) with the complete sample. In the case the complete sample and the subsample share the MRCA they also, with high probability, share the oldest genealogical information. Thus, the effects on the genetic structure of the complete sample of the oldest part of the genealogy are also present in the subsample. In addition, the complete sample and the subsample have had exactly the same timespan to collect mutations. [74] show that the probability that a subsample of a fixed size m shares the MRCA with the complete sample of arbitrarily large size *n* (*n* → ∞) converges to (*m* − 1)/(*m* + 1). Even a subsample of size 2 shares the MRCA with probability 1/3, while a sample of size 19 already shares with probability 0.9. This shows that by this measure (the probability of sharing the MRCA) even a rather small subsample drawn from a large complete sample whose ancestry is governed by the Kingman coalescent captures properties of the complete sample quite well.

The outline of the paper is as follows. In Section 2 we introduce our key object: the probability that the subsample shares the MRCA with the complete sample (see Eq. (3)). In Section 2 we present results for finite sample size, namely Prop. 1 regarding comparing the probability (3) between certain coalescent processes (see Sec. A2 for a precise description of the coalescent processes we consider), Eq. (4) for a recursion to compute (3) exactly for any *Λ*-coalescent (see Eq. (A33)), Prop. 2 which gives a general representation of probability (3) for any *Ξ*-coalescent (see Eq. A32) — and thus for any *Λ* -coalescent — and Prop. 3 which gives a representation of (3) for the Bolthausen-Sznitman coalescent (a specific multiple-merger coalescent, the Beta-coalescent (A36) with *α* = 1). In Section 2.3 we present our main mathematical result (Thm. 1), a representation of the probability (3) as sample size *n* → ∞ for the Beta-coalescent (see Eq. (A36)). We also give a criterion for when the limit of (3), as *n* → ∞, stays positive under a general *Ξ*-coalescent. In Sec. 2.2 we discuss the probability of sharing the oldest allele between the subsample and the larger sample, and the probability of monophyly of the subsample, and we present recursions for these probabilities. In Sec. 3 we investigate by simulations the fraction of internal branch lengths covered by the subsample. Proofs of our mathematical results are presented in Section 4. A brief discussion of the implication of our results, and open problems, is given in Section 5. Section A1 contains a brief description of the population models underlying the coalescent processes we consider, Section A2 contains a detailed description of the coalescent processes, and Section A3 a review of Goldschmidt and Martin’s construction of the Bolthausen-Sznitman *n*-coalescent [39].

For ease of reference we include a table (Table 1) of notation and terminology.

**Table 1.**
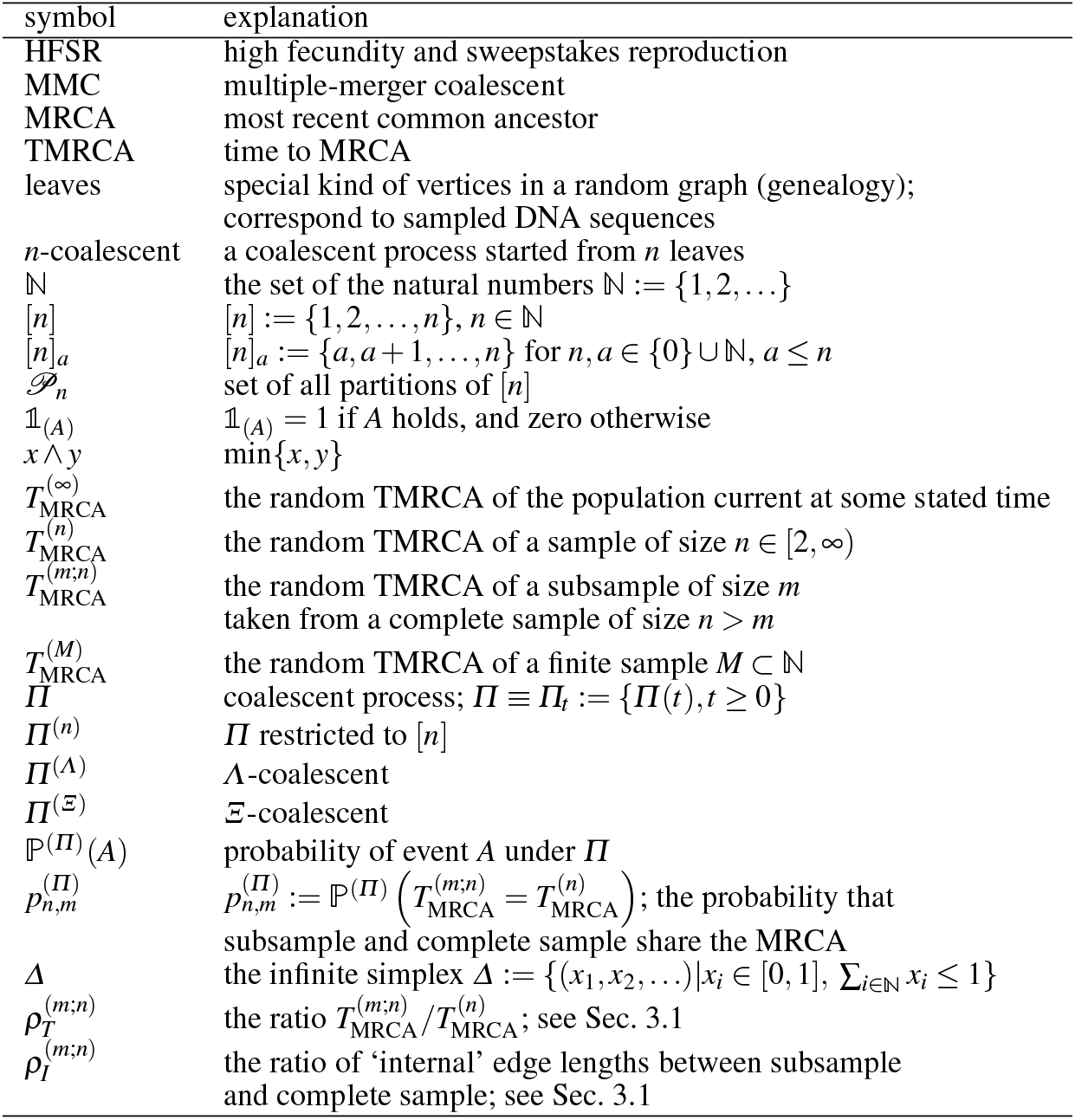
Notation and terminology.

## 2 Sharing the MRCA

We consider a *Ξ*- or *Λ-n*-coalescent with a starting partition *π* = {{1}, …, {*n*}}, i.e. initially all the blocks *π_i_* ∈ *π* are singleton blocks. We refer to the elements of the starting partition as ‘leaves’. A <u>common ancestor</u> of a set 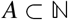 of leaves is any block containing *A*. A set *A* of leaves has a common ancestor if and only if the coalescent passes through a partition with a block containing *A*. This allows us to identify the common ancestor with blocks of the partition-valued states of the coalescent. The MRCA of a set *A* of leaves is the smallest block which contains *A* (whenever that block appears). Given that we start from a <u>finite</u> set [*n*] of leaves (*n* < ∞) we will eventually (i.e. in finite time almost surely) observe the partition {[*n*]} containing only the block [*n*]. Write 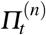 for the partition reached at time *t* in the case when the coalescent process is started from *n* leaves. Let 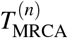 denote the random time to the MRCA (abbreviated TMRCA) of the set [*n*] of leaves, i.e. we define

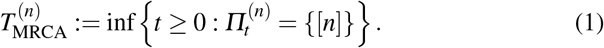

The 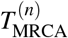 is therefore the first time 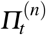 arrives at partition 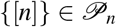, where 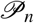 denotes the set of all partitions of [*n*]. Write 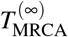 for the TMRCA of the whole population. By a finite <u>sample</u> we mean a finite set A of leaves.

A <u>subsample</u> is a subset of a given sample (a given set of leaves). We let *m* denote the size of the subsample. For convenience and w.l.o.g. we assume leaves 1 to *m* are the leaves of the subsample, and we assume block *π_1_* in any partition always contains element 1. A common ancestor of the subsample is any block containing [*m*]; the MRCA of the subsample is the smallest block containing [*m*] (whenever it appears). We define the TMRCA of a subsample of size m of a sample of size *n* ≤ *m* as

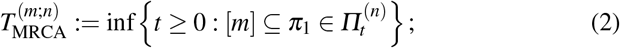

i.e. 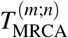 is the time of first occurrence of the subset [*m*] in block *π*_1_ in a partition of *Π*^(*n*)^. The sample and the subsample share the MRCA if the smallest block containing [*m*] ever observed in *Π*^(*n*)^ is [*n*]; this happens almost surely if 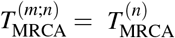.

Our main mathematical results concern the probability

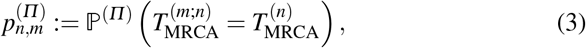

which is the probability that the sample (of size *n*) and the <u>nested subsample</u> (of size *m* < *n*) share their MRCA under the coalescent process *Π*. From now on it should be understood that we always look at nested samples. We are able to obtain representations of 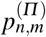 both for finite n and m and also for the limit 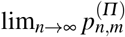, *m* fixed, for some multiple-merger coalescent processes. We will let 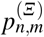 denote 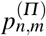 in (3) when *Π* is a *Ξ*-coalescent, and 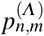 denote 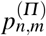 when *Π* is a *Λ*-coalescent.

### 2.1 Finite *n*

Our main focus is to compare genealogical properties of nested samples between different coalescent processes in order to learn what is gained by enlarging the sample size. In this context, a natural question to address is which *n*-coalescent *Π* maximises 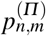 for a given finite sample size n and subsample size m? In the context of *Λ*-coalescents (see Eq. (A33) in Sec. A2) this is the ‘star-shaped’ coalescent with *Λ*-measure *Λ*(*dx*) = *δ*_1_(*x*)*dx* so that *Λ*({1}) = 1, all *n* blocks merge after an exponential waiting time, and 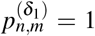. We now compare 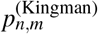 (meaning 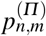 when *Π* is the Kingman-coalescent) to all 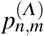 with *Λ*({1}) = 0. We can show the following (see Sec. 4.1 for a proof).

**Proposition 1** *For any given sample size n and subsample size m* < *n there is a Λ′ with Λ′({1}) = 0 which fulfills* 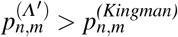.

One can think of *Λ′* as given by *Λ* = *δψ* for some fixed *ψ* ∈ (0, 1) and very close to 1. Prop. 1 holds for any finite sample size *n* and subsample size *m*. Regarding the limit 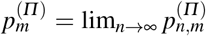 with *m* fixed we conjecture that 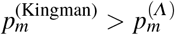 for every *Λ* -coalescent with *Λ* ({1}) = 0. Should our conjecture be true, the limits compare in the opposite way to the comparison of the non-limit probabilities given in Prop. 1.

The result in Prop. 1 holds for a very special *Λ* -coalescent. One can numerically evaluate 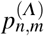 for any *Λ*-coalescent with a recursion (see Sec. 4.7.1 for a proof), and thus compare 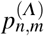 for different *Λ*-coalescents. Let *λ*(*n*) (see Eq. (A34)) denote the total rate of mergers given *n* blocks, and 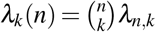 (see Eq. (A33)) denote the rate at which any *k* of *n* blocks merge. Write *β*(*n*, *n* — *k* + 1): = *λ_k_*(*n*)/*λ*(*n*) for the probability of a single merger of *k* blocks (a *k*-merger) given *n* blocks (2 ≤ *k* ≤ *n*). Then

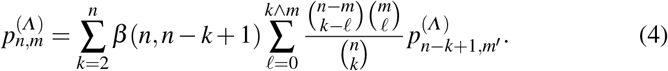

where 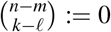 if *n* − *m* < *k* − *ℓ* and 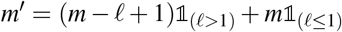. In the case *m* = 2 recursion (4) simplifies to

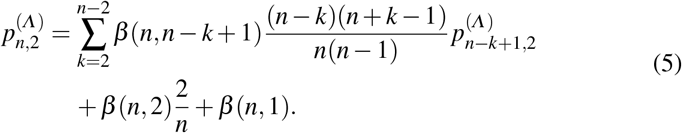

Recursion (4) further simplifies in the case of the Kingman coalescent, since then *β*(*n*, *n* − 1) = 1 for *n* ≥ 2. [74] obtain

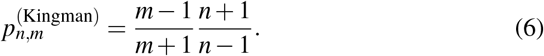

Since the representation (6) only depends on which mergers are possible, the result (6) holds for a time-changed Kingman-coalescent as derived for example in [54] from a population model of ‘modest’ changes in population size.

The Beta-coalescent (see Eq. (A35)) with coalescent parameter *α* ∈ [1, 2), is an example of a *Λ*-coalescent (see Eq. (A33)) and can be derived from population model (A28). Figure 1 shows graphs of 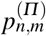 when *Π* is the Beta-coalescent (see Eq. (A35)) as a function of *α*; the results indicate that 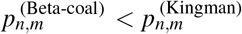 for *n* large enough and any *m*. This shows that one needs a larger subsample under the Beta-coalescent than under the Kingman-coalescent for a given sample size to have the same value of 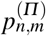. By implication, one gains more information by enlarging the sample under the Beta-coalescent than under the Kingman-coalescent.

**Fig. 1.**
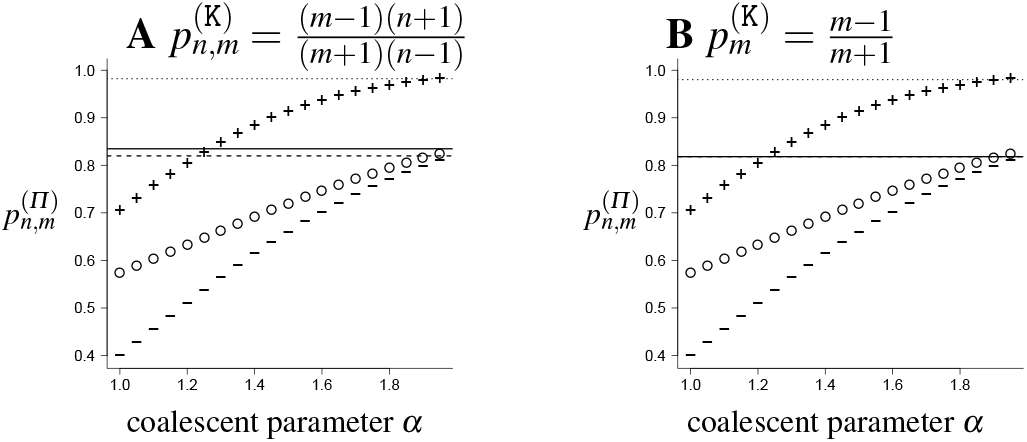
Graphs of 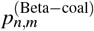 (see Eq. (4)) as a function of *α* for (*n*, *m*) = (10^2^, 10^1^) (circles); (10^3^, 10^1^) (−); (10^3^, 10^2^) (+). The corresponding results for the Kingman-coalescent, 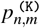 (**A**) and 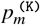 (**B**) are shown as Lines.

We conclude this subsection with two closed-form representations of 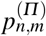. To prepare for the first one we recall the concept of ‘coming down from infinity’.

This property is defined as follows. If a *Ξ-n*-coalescent (see Eq. (A32)) 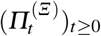 comes down from infinity then, with probability 1, the number of blocks is finite for any *t* > 0, which is equivalent to 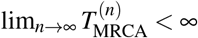 a.s. If 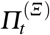, for all *t* > 0, has infinitely many blocks with probability 1, we say that the coalescent ‘stays infinite’. Conditions for *Ξ* to fall into one of these two classes are available, see e.g. [78,77,49]. If 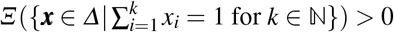, the *Ξ*-coalescent does not stay infinite [77, p.39], but does not necessarily come down from infinity. In fact, there is a.s. a finite (random) time *T* ≥ 0 so that the number of blocks is finite for all *t* > *T* (see [76, p.39]). This means that for such a coalescent, 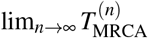 is finite almost surely. For processes that stay infinite 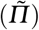, 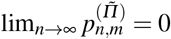 since the MRCA of the set 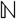 of leaves in the starting partition {{1}, {2}, …} is never reached.

We have a representation of 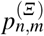 (see Sec. 4.2 for a proof). This representation allows us to later derive characterisations of 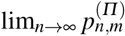 for different multiple-merger coalescents *Π*, see Thm. 1 and Prop. 5. For example, we use Eq. 8 in Prop. 2 to prove Theorem 1.

**Proposition 2** *For any finite measure Ξ on Δ, we have*

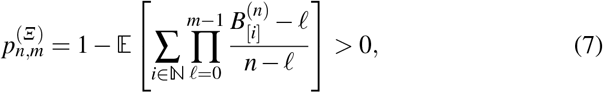

where B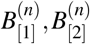, … *are the sizes of the blocks of* 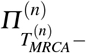, *ordered by size from biggest to smallest where the sequence* 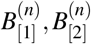, … *is extended to an infinite sequence by taking* 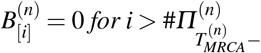. *If the *Ξ*-coalescent comes down from infinity, we have*

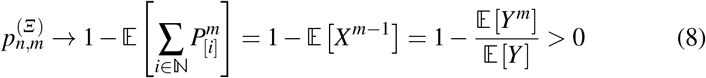

*for fixed m and n* → ∞, where 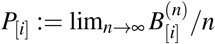 *is the (almost surely existing) asymptotic frequency of the ith biggest block of* 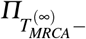, *X is the asymptotic frequency of a size-biased pickfrom the blocks of* 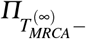, *while Y is the asymptotic frequency of a block picked uniformly at random from* 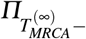.

In the case of the Bolthausen-Sznitman (BS-coal) *n*-coalescent [20], which is a *Λ-n*-coalescent with *Λ*(*dx*) = *dx* (see Eq. (A33)), i.e. the density associated with the uniform distribution on [0, 1], we can give a characterisation of 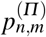 in terms of independent Bernoulli r.v.’s (see Sec. 4.3 for a proof). We use Eq. 9 in Prop. 3 to prove Prop. 5.

**Proposition 3** *Let B*_1_, …, *B_n−1_ be independent Bernoulli random variables with* 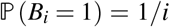. *Let *Π* denote the Bolthausen-Sznitman n-coalescent. For* 2 ≤ *m* ≤ *n*

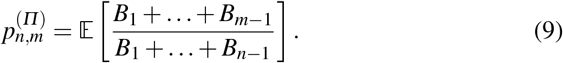

*Moreover*, 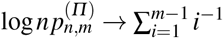 *for n → ∞ and m fixed*.

### 2.2 Two variants of 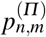

#### 2.2.1 Including the oldest allele

The probability 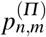 (see Eq. (4)) is an indication of how likely it is that the ‘oldest’ genealogical branches, or the edges connected directly to the MRCA, are (partially) shared between the subsample and the complete sample. We remark that the complete sample and the subsample may share the MRCA without sharing any of the ‘internal’ edges — i.e. an edge subtended by at least 2 leaves (e.g. the marked edges in Fig. 2A) — if the associated coalescent admits multiple mergers (see Fig. 2C for an example). Such events are highly unlikely though for *n* large enough if the *Λ*-coalescent comes down from infinity, see Corollary 1. If the complete sample and the subsample share the MRCA then the subsample is more likely to include the ‘oldest allele’ — i.e. the allele that arose closest to the root — of the complete sample. To derive the actual probability of the event that the subsample carries the oldest allele of the complete sample one needs to include mutation. Consider a *Λ-n*-coalescent with neutral mutation. Mutations are modelled by a homogeneous Poisson point process on the branches of the *Λ-n*-coalescent with (scaled) mutation rate *θ* > 0. We assume the infinitely-many-alleles model. This means that the allelic type of each individual is seen by tracing its ancestral line back to the first mutation on it. The ancestral line shares the type of the MRCA if there is no mutation on the line before the MRCA is reached. We are interested in the event that the oldest allele from the complete sample is also found in the subsample. The probability of this event has been discussed in case of the Kingman’s *n*-coalescent [74] (see Eq. 5.13). For multiple-merger coalescents this probability can be expressed by using the concept of ‘frozen’ and ‘active’ ancestral lines in a *n*-coalescent with mutation [23]. At a given time *t*, an ancestral lineage is called frozen if there has been a mutation on it, otherwise it is called active. The age of a sampled allele (*i*, say) is the waiting time *τ_i_* until its’ ancestral lineage is frozen. For consistency we prolong the *n*-coalescent after reaching the MRCA (at time 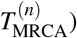) by a single ancestral line. The first mutation on the prolonged line is seen after an additional Exp(*θ*/2) time which freezes the line. Thus, the oldest allele of a sample is given by the ancestral lineage which is frozen last (active the longest), and this age is max{*τ_i_*: *i* ∈ [*n*]} for the sample and max{*τ_i_*: *i* ∈ [*m*]} for the subsample. Let *A^(n)^*(*t*) denote the count of active ancestral lineages in the sample at time *t*. We write

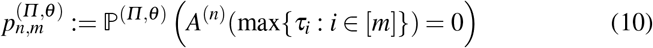

**Fig. 2.**
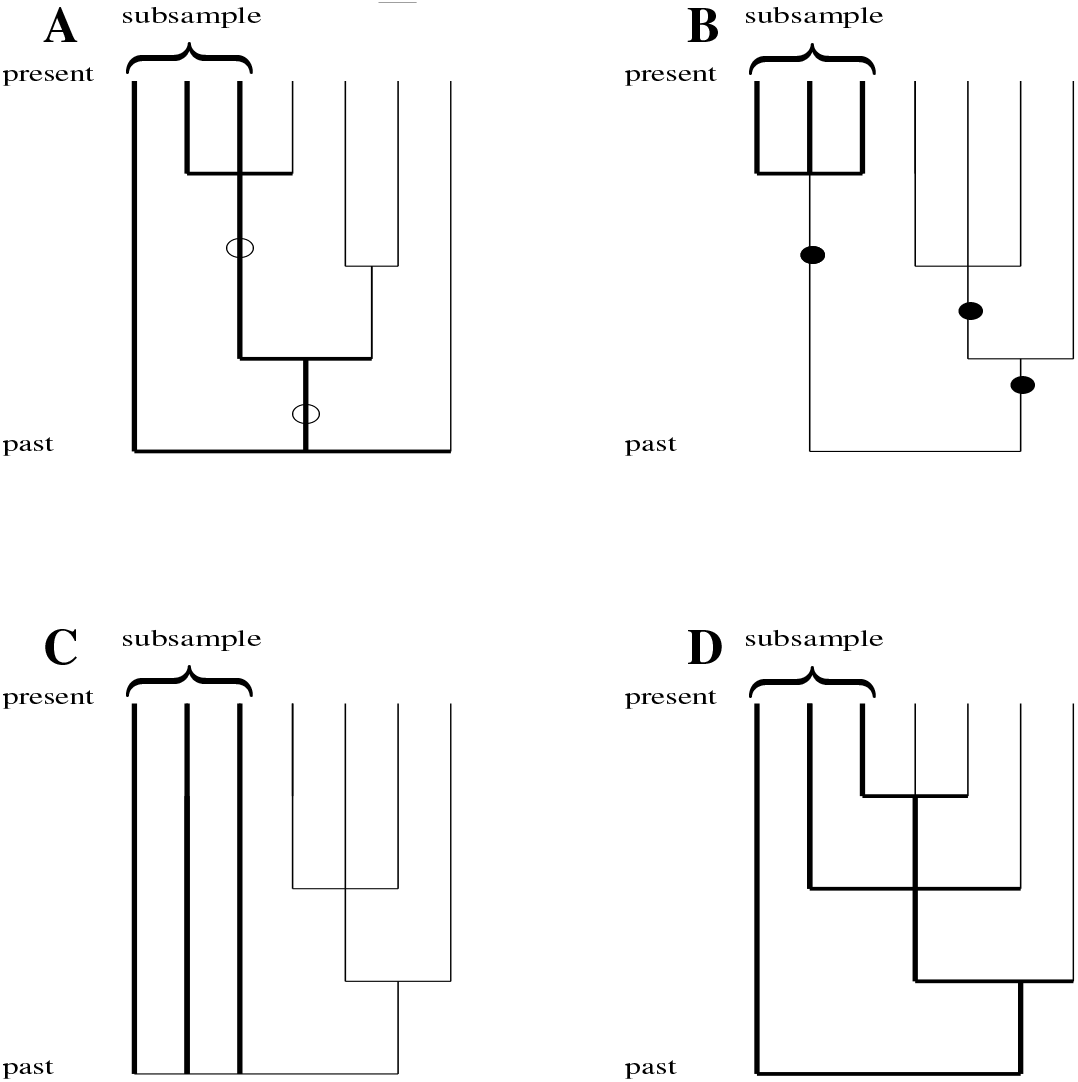
Examples of genealogies. Thick edges denote lineages ancestral to the subsample of size m = 3; sample size *n* = 7. The marked edges in **A** denote internal ancestral lineages to both the subsample and the complete sample; the marked edges in **B** denote lineages internal only to the complete sample. In **C** the complete sample and subsample share the MRCA without sharing any of the internal edges. The genealogies are shown from the time of sampling (present) until the MRCA of the complete sample is reached (past). In **C** the complete sample and the subsample share the MRCA without sharing any internal edges. In **D** the complete sample and the subsample share the MRCA and <u>all</u> the internal edges.

for the probability that the subsample includes the oldest allele of the sample.

We consider 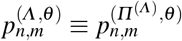 for 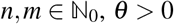. The case *n* = *m* = 1 (or *n* > *m* = 1) means we trace back a single lineage until it is hit by a mutation (either in the sample and/or subsample). The boundary conditions are 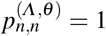 and 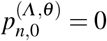 for *n* > 0. The recursion for 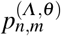 is

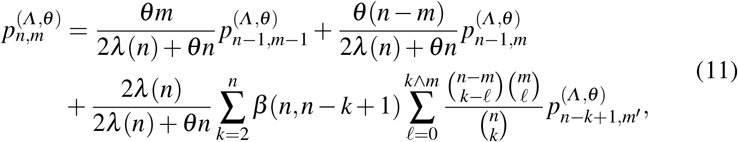

where 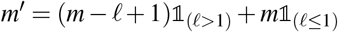 (see Sec. 4.7.3 for a proof),

The probability 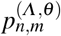 is a function of the scaled mutation rate *θ*. Here, and in most models in population genetics which include mutation, *θ*: = *μ_N_*/*c_N_* where *μ_N_* is the rate of mutation per locus per generation, and c_N_ is the pairwise <u>coalescence probability</u>, or the probability that 2 distinct individuals sampled at the same time from a population of size *N* have the same parent. Since (usually) one arranges things so that *c_N_* → 0 as *N* → ∞ to ensure convergence to a continuous-time limit [71,64], and since *θ* is usually assumed to be of order 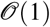, we let *μ_N_* depend on *N*. The key point here is that *θ* depends on *c_N_*. By way of an example, *c_N_* = 1/*N* for the haploid Wright-Fisher model, while 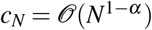 for the Beta(2 − *α*, *α*)-coalescent, 1 < *α* < 2 [72]. This means that the scaled mutation rates (*θ*) are not directly comparable between different coalescent processes; this again means that expressions 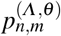, defined in (10), for example) which depend on the mutation rate cannot be directly compared between different coalescent processes that may have different timescales. We further remark that we must define *θ* to be proportional to 1/*c_N_* since the branch lengths on which the mutation process runs are in units of 1/*c_N_* (i.e. 1 coalescent time unit corresponds to ⌊1/*c_N_*⌋ generations); thus if we don’t rescale the mutation rate *μ_N_* with 1/*c_N_* we would never see any mutations. It is therefore the mutation rate *μ_N_*, which must be determined from molecular (or DNA sequence) data, which determines the timescale; the quantity *c_N_* comes from the model.

#### 2.2.2 The smallest block containing [*m*]

The probability 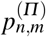 is also the probability of the event that the MRCA of the subsample (of size *m*) subtends all the *n* leaves ([*n*] is the smallest block containing [*m*]). A related more general question is to ask about the distribution of the size (number of elements) of the smallest block which contains [*m*]. This is the same as asking about the distribution of the number of leaves subtended by the MRCA of the subsample. For Kingman’s *n*-coalescent, the distribution is computed in [82, Thm. 1]. The probability of the event that the MRCA of the subsample subtends only the leaves of the subsample is especially interesting, see e.g. [88, p. 184, Eq. 2], where this probability is described recursively in the case of the Kingman-coalescent. This recursion can be easily extended to *Λ*-coalescents. Define 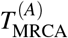 to be the first time that A is completely contained within a block of *Π_t_*. Write

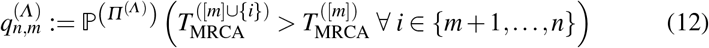

for the probability that the MRCA of the subsample subtends only the leaves of the subsample (leaves 1 to *m*). Let *β*(*n*, *n* − *k* + 1) be the probability of a *k*-merger (2 ≤ *k* ≤ *n*) given *n* active lines. The recursion for 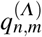 is (see Sec. 4.7.2 for a proof)

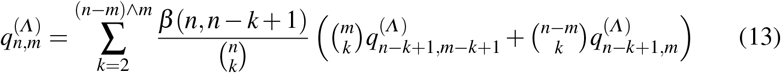

with boundary conditions 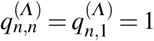 for 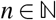. One may use 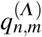 to calculate the *p*-value of a test for monophyly or non-random mating (see Discussion in [82]), i.e. to calculate the *p*-value of a test for observing block [*m*] under the null-hypothesis that the *Λ*-coalescent models the genealogy.

As one might expect (see Sec. 4.3 for a proof), for *m* fixed and for any *Λ*-coalescent,

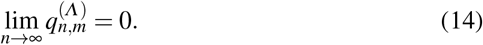

In the case of the Bolthausen-Sznitman (BS-coal) *n*-coalescent we obtain an exact representation of 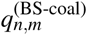 (see Sec. 4.4 for a proof).

**Proposition 4** *Let* 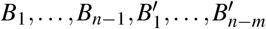 *be independent Bernoulli variables with* 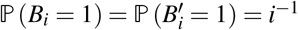. *For the Bolthausen-Sznitman *n*-coalescent we have, for* 2 ≤ *m* < *n*,

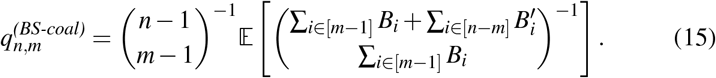

### 2.3 The limit 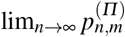

As we stated in the Introduction the aim of modelling the random genealogy of a sample of DNA sequences drawn from some population is to learn about the evolutionary history of the population. We are therefore interested in investigating the behaviour of the genealogical statistics within our framework of nested samples as the size of the complete sample is allowed to be arbitrarily large, but keeping the size of the subsample fixed. In this subsection we discuss the limit 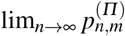 with *m* fixed. For a fixed 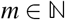, write

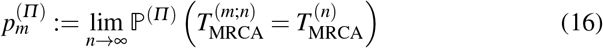

for the probability, under coalescent *Π*, that a subsample of size *m* shares the MRCA with an arbitrarily large sample. The limit 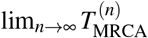 is a valid limit for any coalescent (even if it diverges) and therefore (16) is well defined. For any *Ξ*-coalescent 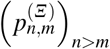 is monotonically decreasing as *n* increases. The limit 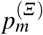 is derived under the assumption that the same *Ξ*-coalescent is obtained for arbitrarily large sample size. This assumption may not hold when one wants to relate to finite real populations. The quantity pshould therefore only be regarded as a limit. See further discussion on this point in Sec. 5.

For the Kingman-coalescent we have the following result, first obtained in [74] by solving a recursion,

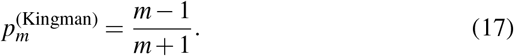

To see (17) without solving a recursion, we consider the process forwards in time from the MRCA. Label the two ancestral lines generated by the first split (of the MRCA) as *a*_1_ and *a*_2_. The fraction of the population that is a descendant of *a*_1_ is distributed as a uniform random variable on the unit interval, see e.g. the remark after Thm. 1.2 in [8]. Therefore, with *U* a uniform r.v. on [0, 1], and any finite 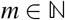,

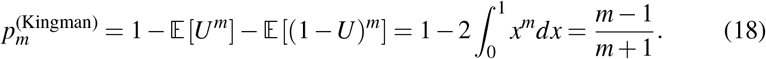

For the Bolthausen-Sznitman coalescent (BS-coal) 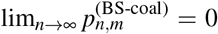 for *m* fixed, see Prop. 3. We remark in this context that the Bolthausen-Sznitman coalescent does not come down from infinity.

Result (17) indicates that 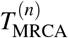 is a good statistic for capturing a property of the population with a small sample, at least under the Kingman coalescent. We remark that the Kingman coalescent comes down from infinity. Result (17) (and (18)) is the ‘spark’ for the current work.

Our main mathematical result, Thm. 1, is a representation of 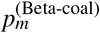, i.e. 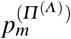 is the Beta(2 − *α*, *α*)-coalescent [79] (Beta-coalescent; see Eq. (A36)). For *α* ∈ (1, 2), the Beta-coalescent comes down from infinity. The representation of 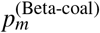 given in Thm. 1 can be directly derived from [8, Thm. 1.2], which is a result based on the connection between the Beta-coalescent and a continuous-state branching process (see Sec. 4.5 for a proof).

**Theorem 1** *Define* 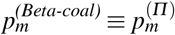 *(see Eq. (16)) when Π is the Beta-coalescent for α* ∈ (1, 2). *Let K denote the random number of blocks involved in the merger upon which the MRCA of* [*n*] *is reached; K has generating function* 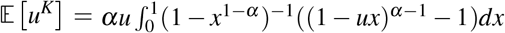 *for u* ∈ [0, 1] *[48, Thm. 3.5]. Let* 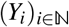 *be a sequence of i.i.d. r.v. with Slack’s distribution on* [0, ∞), *i.e. Y*_1_ *has Laplace transform* 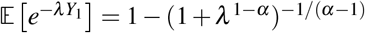 *[81]. We have the representation*

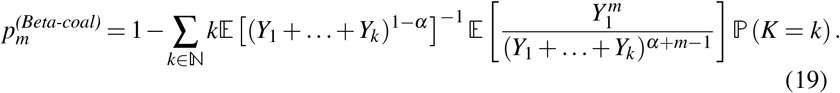

We discuss the relevance of 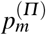 for biology in Sec. 5.

We close this subsection with a consideration of the limit 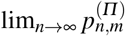 when *Π* is a *Ξ*-coalescent (A32). We give a criterion for when 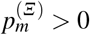. This question is closely related to the question of coming down from infinity for a *Ξ*-coalescent. We have the following result (see Sec. 4.6 for a proof).

**Proposition 5** *Consider any Ξ-coalescent. For any fixed* 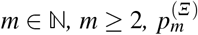 *exists. If the coalescent comes downfrom infinity or* 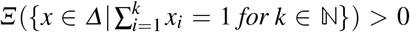 then 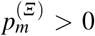. If it stays infinite then 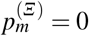.

## 3 Relative times and lengths

In this section we use simulations to assess how well the subsample’s genealogy ‘covers’ the genealogy of the complete sample containing the subsample.

We consider the relative times 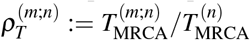 and the relative lengths 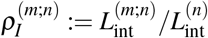 where 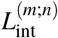 is the sum of the lengths of the ‘internal’ edges associated with the subsample and 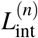 is the sum of the lengths of internal edges of the complete sample. An edge (ancestral line) is internal if it is subtended by at least two leaves, else it is ‘external’. An edge is associated with the subsample if at least one of the leaves subtending it belongs to the subsample — we call such a line a subsample line. By way of example, the continuing line of the first merger in Fig. 2D counts as an internal subsample line, although it is only subtended by a single leaf of the subsample. The ratio 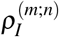 keeps track of the fraction of internal edges of the sample’s genealogy that are covered by edges of the subsample’s genealogy. The statistic 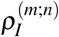 indicates how much of the ‘ancestral variation’, or mutations present in at least 2 copies in the sample, are captured by the subsample. The ratio 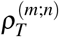 indicates how likely we are to capture with the subsample the ancestral variation in the complete sample. We compare 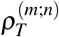 and 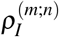 between the Beta-coalescent, a time-changed Kingman coalescent representing Wright-Fisher (or a similar) sampling with exponential population growth (see Sec. A1), and the classical Kingman-coalescent.

### 3.1 Simulation method

We simulate realisations of 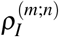 and 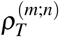 for the classical and time-changed Kingman-coalescents and for Beta-coalescents. All processes have a Markovian jump chain and waiting times between the jumps are dependent on the current state of the jump chain, more precisely on the number of ancestral lines present. We simulate sample genealogies for a sample of size *n* by first generating the jump chain, i.e. choosing how many ancestral lines are merged. Given the size of a merger we draw the number of internal and external lines to merge. Given a number of sample lines we draw the waiting time until the next merger.

Let *j* denote the current number of lines of the complete sample. Let *M* ∈ {2, …, *n*} be the size of the next merger. Under a (time-changed) Kingman-*n*-coalescent *M* = 2 regardles of the value of *j* ≥ 2. Under a *Λ-n*-coalescent *M* = *k* with probability 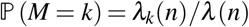 (see Eq. (A33) and (A34)) for 2 ≤ *k* ≤ *n*. The *k* lines picked are merged into a single ancestral line, so *j* − *k* + 1 lines are left after the merger. We draw subsequent mergers starting with *n* lines until there is only a single line left, when the MRCA of the sample is reached.

Consider 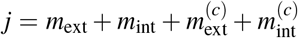 sample ancestral lines present, from which *m*_ext_, *m*_int_ are external and internal subsample lines, whereas 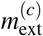 and 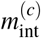 are external and internal lines not subtended by leaves from the subsample. As an example, we start (before any merger) with *n* lines distributed as 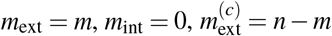 and 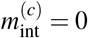. All *n*-coalescents are exchangeable, and we always pick lines to merge at random from the lines present without replacement. This leads to drawing numbers of lines *x*_1_, *x*_2_, *x*_3_, and *x*_4_ from the four categories *m*_ext_, *m*_int_, 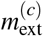 and 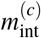 following a multivariate hypergeometric distribution (<u>*X*</u> = (*X*_1_, …, *X*_4_))

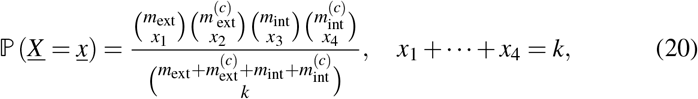

where *k* denotes the given merger size. All lines are merged into a single ancestral line, which is a subsample line if and only if at least one subsample line was picked in the merger (*x*_1_ + *x*_3_ ≥ 1), so the numbers of lines belonging to the four categories change from before to after the merger as

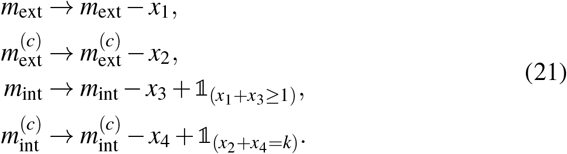

The transitions shown in Eq. (21) reflect our assumption that if at least 1 subsample line is involved in a given merger, the continuing ancestral line is considered to belong to the subsample; mutations that arise on the continuing line will then be carried by the subsample, and visible in the subsample unless all the subsample lines were involved in the merger (*x*_1_ = *m*_ext_ and *x*_3_ = *m*_int_ and *x*_1_ + *x*_3_ ≥ 1). Therefore, if a single external subsample line, and no other subsample line, is involved in a merger (*x*_1_ = 1, *x*_3_ = 0) we regard the continuing line as an ‘internal’ line of the subsample. An external line of the subsample therefore remains so only until it is involved in a merger. By way of example, the continuing line of the first merger in Fig. 2D counts as an internal line of the subsample. The continuing line of the first merger in 2B is <u>not</u> a subsample line since the MRCA of the subsample leaves is reached in the first merger; the continuing line of the first merger in 2B counts as an internal line of the complete sample.

Denote by *T_j_* the random waiting time for the first merger of the *j*-coalescent. The coalescent process under exponential population growth is a time-changed Kingman-coalescent (see e.g. [25,30]). [41] give a way of sampling *T_j_* under exponential growth. Let *β* > 0 denote the growth rate under exponential growth.

Write *S_j_* = *T_n_* + ⋯ + *T_j_* for 2 ≤ j ≤ *n*, with *S*_*n*+1_ = 0 a.s. If {*U_j_*: 2 ≤ *j* ≤ *n*} denotes a collection of i.i.d. uniform (0,1] random variables, then [41]

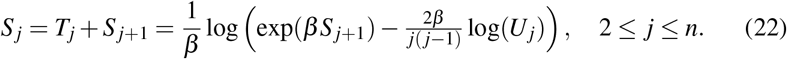

Eq. (22) tells us that if *β* is very large, the time intervals *T_j_* near the MRCA become quite small. The time intervals near the leaves are much less affected. We choose the grid of values for *β* as

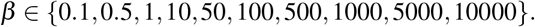

Recall in this context the growth model *N_k_* = *N*_0_(1 + *β/N*_0_)*^k^* for the population size in generation *k* ≥ 0 going forward in time, and where *N*_0_ is the population size at the start of the growth (*k* = 0). Our choice of grid values for β should reflect the range of growth from weak (*β* = 0.1) to very strong (*β* = 10^4^) and most estimates of *β* obtained for natural populations should fall within this range.

Under the Beta-coalescent without growth *T_j_* is an exponential with rate *λ*(*j*) = *λ*_2_(*j*) + ⋯ + *λ_j_*(*j*) where *λ_i_*(*j*) is given in Eq. (A36).

A realisation of 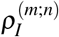 is obtained as follows. Given 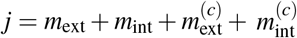 current sample lines, let *t_j_* denote a realisation of *T_j_*, the random time during which there are *j* lines of the complete sample. We update the total lengths 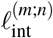 of internal subsample lines, and 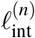 of internal lines of the complete sample, as

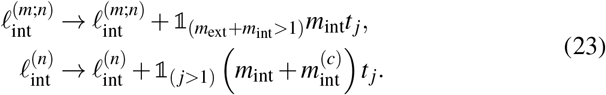

The updating rule for 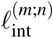 in Eq. (23) reflects the fact that mutations on the common ancestor line of the subsample, for example the continuing line after the merger of all 3 subsample lines in Fig. 2D, are not visible in the subsample. The updating rule for 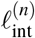 in Eq. (23) similarly reflects the fact that mutations on the continuing line of the MRCA of the complete sample are not visible in the sample; but once the MRCA of the complete sample is reached we stop the process.

A realisation of 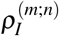 is then recorded as 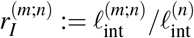. By way of example, the edges marked with a black dot in Fig. 2B are internal edges of the complete sample while the edges marked with a circle in Fig. 2A are internal edges associated with the subsample as well as the complete sample, and we have 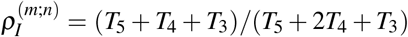 for the genealogy in Fig. 2A. There are no internal edges associated with the subsample in Fig. 2B and 2C; therefore 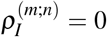 for the genealogies in Fig. 2B and 2C. The sample and the subsample share <u>all</u> the internal edges in the genealogy shown in Fig. 2D and therefore 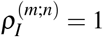.

Realisations of 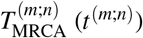 and 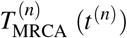 are recorded as

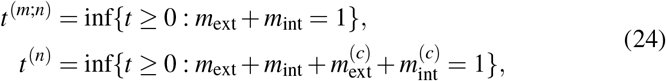

by adding up the realised waiting times *t_j_* of *T_j_*. We record a realisation of 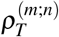 as 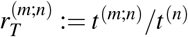.

### 3.2 Simulation results

Figures 3 and 4 show estimates, in the form of violin plots[44], of the distributions of 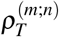 (left column) and 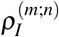 (right column); under the Beta-coalescent as a function of *α* (Figure 3) and under exponential growth as a function of *β* (Figure 4). We see that under exponential growth the distribution of pT ‘ can be rather concentrated (recall Eq. (22)). The estimates shown in Fig. 4 of the distribution of 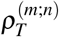 indicate that 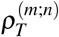 becomes more concentrated at 1 as β increases. Recall in this context that 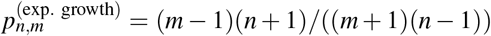 since exp. growth results in a time-changed Kingman-coalescent.

**Fig. 3.**
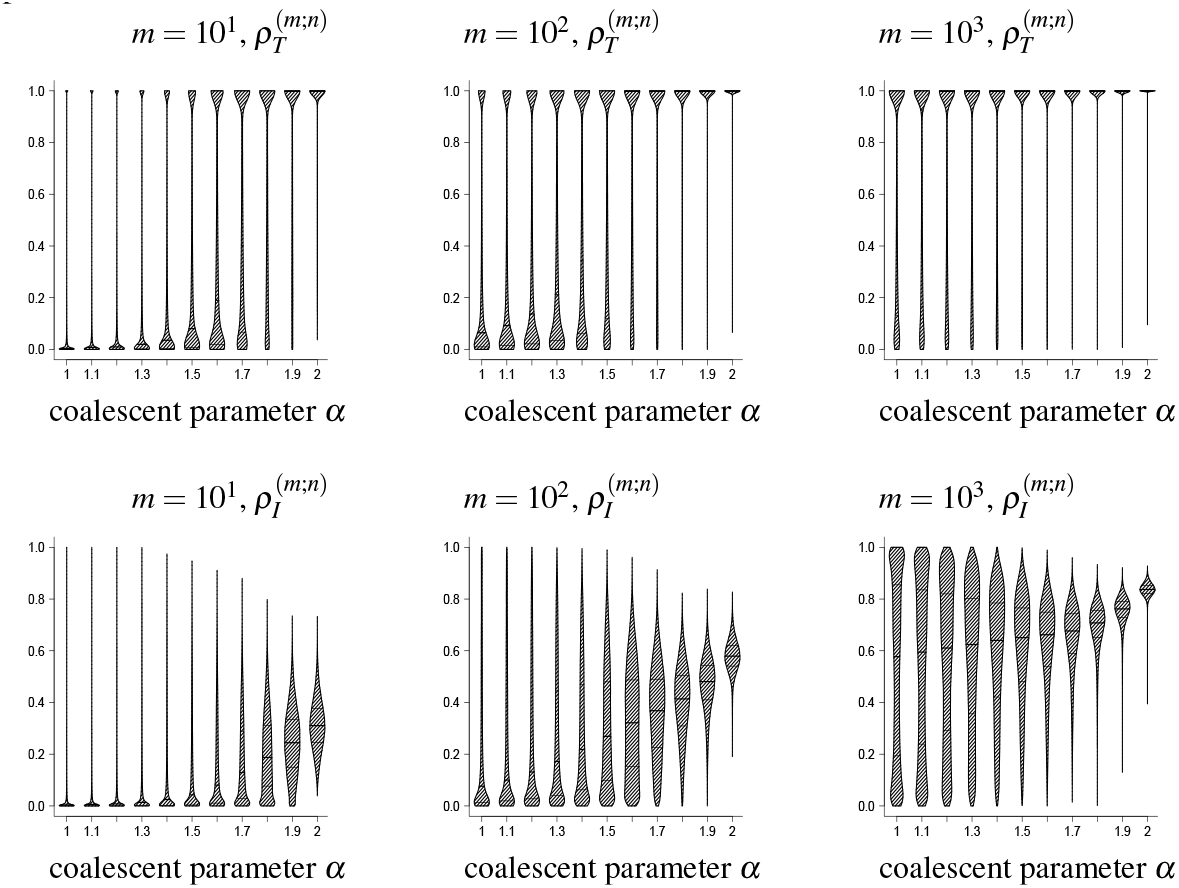
Estimates, shown in the form of violin plots [44] of the distributions of 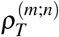 and of 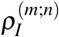 as functions of the coalescent parameter *α* of the Beta(2 − *α*, *α*)-coalescent for values of sample size *n* = 10^4^ and subsample size *m* as shown. The coalescent process at *α* = 2 is the Kingman-coalescent. For explanation of symbols see Subsection 3.1. Shown are results from 10^5^ replicates.

In Figure 3 we see a gradual shift in the distribution of 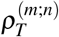 as subsample size increases; from being skewed to the right (ie. towards higher values) to being skewed to the left (ie. towards smaller values). This is in sharp contrast to the distribution under exponential growth (Figure 4) where the distribution of 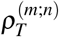 is always skewed to the left. This indicates that under a multiple-merger coalescent process a subsample is much less informative about the complete sample than under exponential growth. In contrast, under exponential growth, even a small subsample can be very informative about the complete sample, especially in a strongly growing (large *β*) population. Estimates of the means 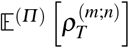, shown in Figure 5 (circles) for the Beta-coalescent, and in Figure 6 (circles) for exponential growth, further strengthen our conclusion.

**Fig. 4.**
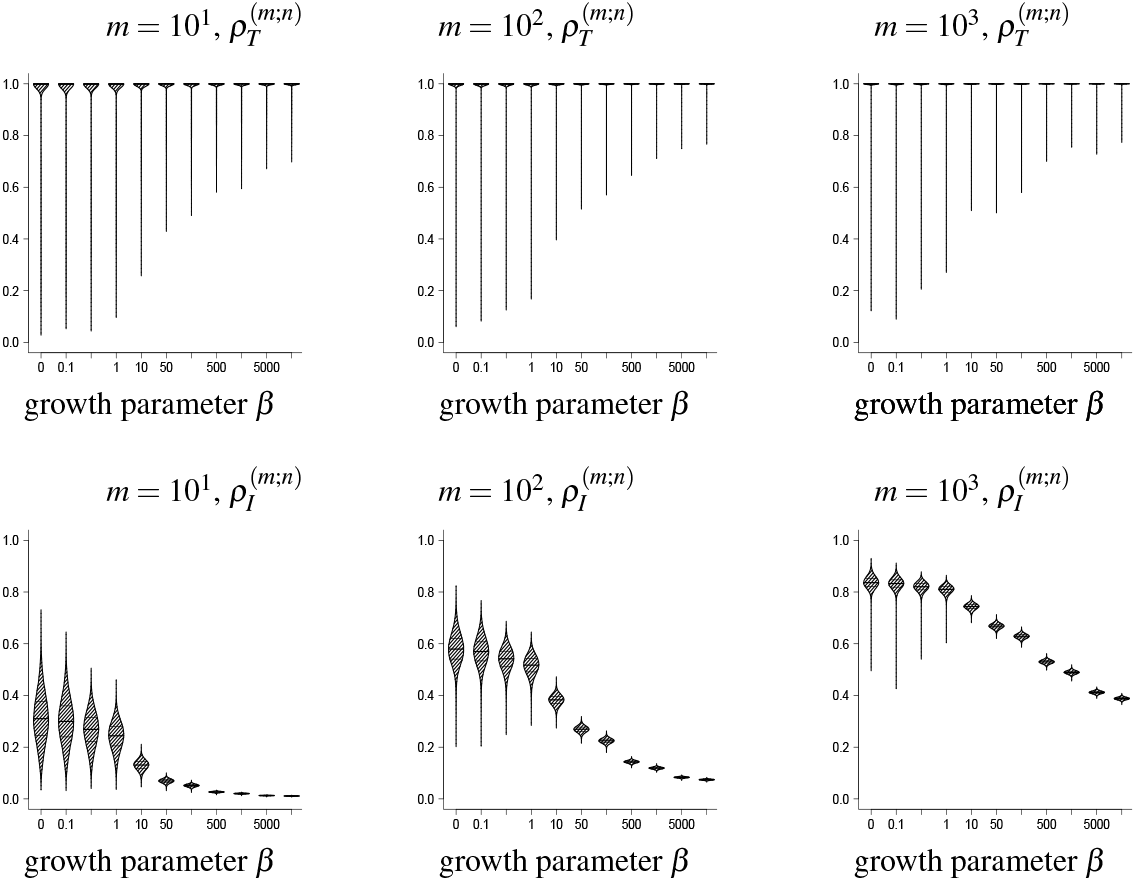
Estimates, shown in the form of violin plots [44] of the distributions of 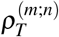 and of 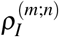 as functions of the exponential growth parameter *β* for values of sample size *n* = 10^4^ and subsample size m as shown. The coalescent process at *β* = 0 is the Kingman-coalescent. For explanation of symbols see Subsection 3.1. The grid of values of *β* is {0.1, 0.5, 1.0, 10.0, 50.0, 100.0, 500.0, 1000.0, 5000.0, 10000.0}. Shown are results from 10^5^ replicates.

**Fig. 5.**
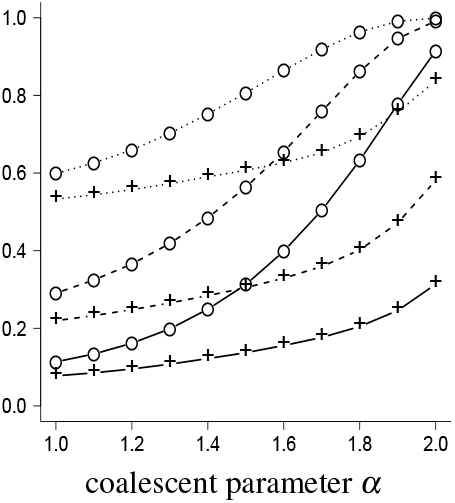
Estimates of 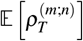 (∘), and of 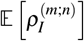 (+) as functions of the coalescent parameter *α* for values of sample size *n* = 10^4^ and subsample size *m* = 10^1^ (solid Lines); *m* = 10^2^ (dashed Lines); *m* = 10^3^ (dotted lines). The coalescent process at *α* = 2 is the Kingman-coalescent. For explanation of symbols see Subsection 3.1. Shown are results from 10^5^ replicates.

**Fig. 6.**
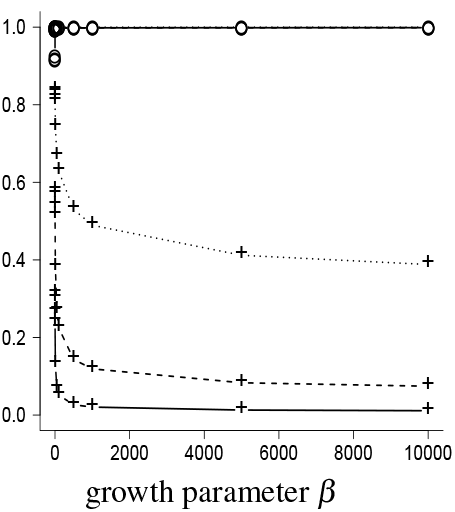
Estimates of 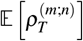 (∘) and of 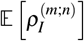 (+) as functions of the exponential growth parameter *β* for values of sample size *n* = 10^4^ and subsample size m = 10^1^ (solid lines); m = 10^2^ (dashed lines); m = 10^3^ (dotted lines). The coalescent process at *β* = 0 is the Kingman-coalescent. For explanation of symbols see Subsection 3.1. The grid of values of *β* is {0.1, 0.5, 1.0, 10.0, 50.0, 100.0, 500.0, 1000.0, 5000.0, 10000.0}. Shown are results from 10^5^ replicates.

The distribution of 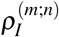, the relative lengths of internal edges, also behaves differently between the Beta-coalescent and exponential growth. The distribution of 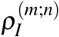 becomes more concentrated around smaller values as growth becomes stronger (*β* increases) while it stays highly variable as *α* tends to 1, although the median decreases as skewness increases (*α* tends to 1). This indicates that we capture less and less of the ‘ancestral variation’ (mutations observed in at least 2 copies in the sample) in the complete sample as growth or skewness increase. Estimates of 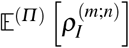 (Figures 5 and 6, ‘+’) also indicate that one would need a large sample to capture at least half of the ancestral variation if growth or skewness is high.

To conclude, 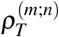 and 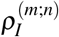 seem to tend to opposite values under exponential growth; 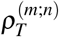 to 1 and 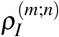 to small values, as *β* increases. Thus, even if we are sharing the MRCA with higher probability as *β* increases (recall that the samples are nested), we are capturing less and less of the ancestral variation. Essentially the opposite trend is seen for both 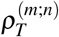 and 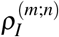 under the Beta-coalescent; the distributions of both statistics stay highly variable as *α* → 1.

## 4 Proofs

### 4.1 Proof of Prop. 1

#### Proof

Let *Λ* = *δ_p_* for *p* ∈ (0, 1) which fulfills *Λ*({0}) = 0. Clearly

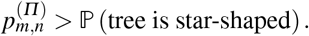

The probability that the associated *Λ-n*-coalescent is star-shaped, i.e. all blocks merge at the first (and then only) collision, is

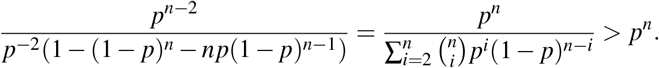

For any star-shaped path of a *n*-coalescent, we have 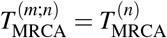 for any *m* < *n*. Thus, we can choose *Λ′* s.t. *Λ′* = *δ_p_* with

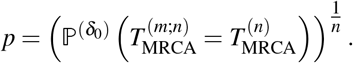

### 4.2 Proof of Prop. 2

#### Proof

Assume *Π* is a *Ξ*-coalescent. The event 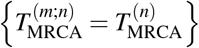 is the complement of the event

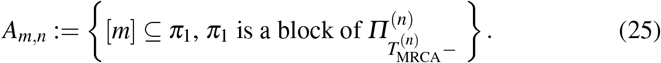

Due to the exchangeability of the *Ξ*-coalescent, 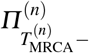 is an exchangeable partition of [*n*]. Given the (ordered) block sizes 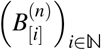, the probability that a given block of size 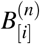 contains [*m*] is given by drawing without replacement, i.e.

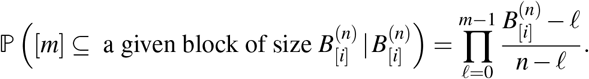

Summing this up over all blocks and taking the expectation yields 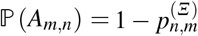, thus establishing Eq. (7) (by definition there is more than one block at time 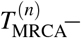, i.e. at time infinitely close to 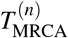, so 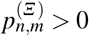.)

To show the convergence in Eq. (8) we first establish that all objects are well defined. Assume now that the *Ξ*-coalescent comes down from infinity, so at any time *t* > 0, there are only finitely many blocks in the partition *Π_t_* almost surely. For *n* → ∞, Kingman’s correspondence [56, Thm. 2] ensures that the asymptotic frequencies of the blocks in the partition *Π_t_* of 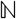 exist almost surely and are limits of the block frequencies in the *n*-coalescent as written in the proposition. Pick an arbitrarily small *t* > 0. Then, consider only paths where *Π_t_* has more than one block. Since the number of blocks of *Π_t_* is finite a.s. we can find 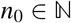 so that 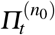 has at least one individual in any block of *Π_t_* (thus has the same number of blocks). By construction, from time t onwards, the *Ξ-n*_0_-coalescent merges the blocks in exactly the same (Markovian) manner as the *Ξ*-coalescent. So if *Π_t_* has more than one block, 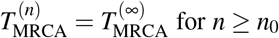 for *n* > *n*_0_ and the asymptotic frequencies at 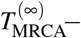 exist (since their corresponding blocks are a specific merger of the blocks of *Π_t_* whose block frequencies exist). Now 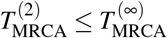 almost surely and 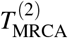 is Exp(*Ξ*(Δ))-distributed. Therefore, for almost every path, we can choose 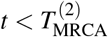 so that *Π_t_* has more than one block.

We have established that all objects are well defined; now we show the actual convergence in (8). For ***x*** ∈ Δ let

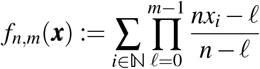

and 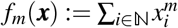. We have *f_n,m_* → *f_m_* uniformly on Δ and that *f_m_* is continuous on Δ in the ℓ^1^-norm with 0 ≤ *f_m_* ≤ 1. We can rewrite, using Eq. (7),

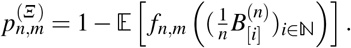

For any *ε* > 0 we find *n*_0_ so that for *n* ≥ *n*_0_

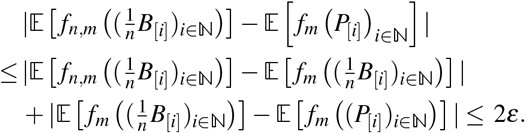

We have used uniform convergence of *f_n,m_* to *f_m_* to control the first difference and the convergence (in law) of 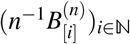 to 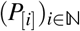 to control the second.

The representation of the limit in Eq. (8) in terms of *X* and *Y* follows directly from the properties of exchangeable partitions (c.f. for example [9,10]). The first equality is [9, Eq. (1.4)], while the second equality uses the correspondence between the distribution of a size-biased and a uniform pick of a block, see [9, Eq. (1.2)]. By definition 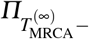 has more than one block almost surely so the limit in Eq. (8) is > 0.

*Remark 1* Reordering the block frequencies, e.g. in order of least elements of blocks, does not change Eq. (8).

### 4.3 Proof of Prop. 3

#### Proof

We use the construction of [39] in which the Bolthausen-Sznitman coalescent is obtained by cutting a random recursive tree 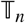 with *n* nodes at independent Exp(1) times, see Sec. A3. Consider the last merger in the Bolthausen-Sznitman *n*-coalescent. In terms of cutting edges of 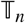, the last merger is reached when the last edge connected to the root of 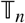 is cut. Let *E_n_* be the number of such edges in 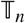. For 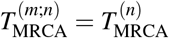, we need that not all *i* ∈ [*m*] are in a single block of the *n*-coalescent before the last merger (see proof of Prop. 2).

By construction, for any node with label in [*m*], on the path to the node labelled 1 (root) in the uncut tree 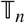, the last node passed before reaching the root must also have a label from [*m*]. Thus, any node connected to the root of 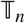 that is labelled from [*n*]_*m*+1_ cannot root a subtree that includes any nodes labelled from
[*m*].

Now, we consider the last edge of 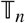 cut in the construction of the Bolthausen-Sznitman *n*-coalescent, which causes the MRCA of the *n*-coalescent to be reached. It has to be connected to the root. Consider the two subtrees on both sides of the edge cut last. One subtree contains the root, thus includes at least the label 1 from [*m*]. If the other subtree is rooted in a node labelled from [*m*], we have 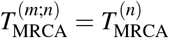, since both subtrees contain labels of [*m*], thus not all *i* ∈ [*m*] are in a single block of the *n*-coalescent before the last merger. If the subtree not containing the root has a root labelled from [*n*]_*m*+1_, as argued above, it contains no labels from [*m*]. Additionally, since we are at the last cut, all other edges connected to the root of 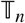 have already been cut and all labels in the subtrees rooted by them joined with label 1. Thus, all labels in [*m*] are labelling the root before the last cut, which corresponds to [*m*] being a subset of a block of the *n*-coalescent before the last merger hence 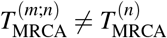.

This shows 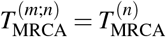 if and only if the last edge cut is an edge connecting a node labelled from [*m*] with the root. Let *E_m_* be the count of edges of 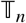 connected to the root labelled from [*m*] and *E_n_* be the total count of edges connected to the root. Then,

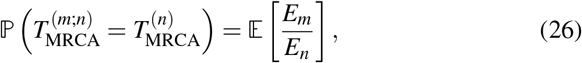

because given 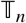, *E_m_*/*E_n_* is the probability that the edge cut last is connected to a node with a label from [*m*]; edges are cut at i.i.d. times, so the edge cut last is uniformly distributed among all edges connected to the root.

As we see from the sequential construction of 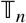, *E_m_* is the number of edges connected to 1 when the first m nodes are set, the resulting tree is a random recursive tree 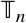 with *n* leaves. The numbers *E_n_* and *E_m_* can be described in terms of a Chinese restaurant process (CRP), see [39, p. 724]: The number of edges connected to node 1 is distributed as the number of tables in a CRP with *n* (resp. *m*) customers.

This distribution is 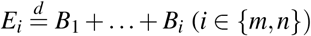, where *B*_1_, …, *B*_2_ are independent Bernoulli variables with *P*(*B_j_* = 1) = *j*^−1^, see e.g [3, p. 10]. The sequential construction of the random recursive trees (and the connected CRPs) ensures that the *B*_1_, …, *B_m_* are identical for *E_m_* and *E_n_*. This establishes the equality of Equations (26) and (9).

From the proof of [38, Lemma 3], we have 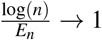 in *L*^1^ for *n* → ∞. The sequence 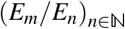 is bounded a.s. Thus, bounded convergence ensures

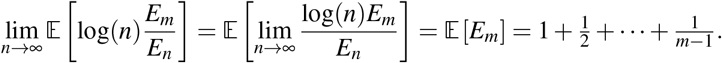

### 4.4 Proof of Prop. 4

#### Proof

As in Subsection 3 we use the construction of the Bolthausen-Sznitman *n*-coalescent described in [39]. We wish to establish the probability that the MRCA of a subsample of size *m* from a sample of size *n* is an ancestor of only the subsample in the *n*-coalescent. We will also use the Bernoulli variables *B_i_*, *i* ∈ [*n*] of 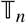 as in Prop. 3, where *B_i_* = 1 if the node labelled *i* is directly connected to the root (node labelled 1). If we look at the cutting procedure which constructs the Bolthausen-Sznitman *n*-coalescent from 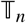, we observe that no path of 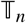 can contribute positive probability to 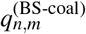 that attaches any node labelled from [*n*]_*m*+1_ to a node labelled from [*m*]_2_. If we do attach a node labelled *i* ∈ [*n*]_*m*+1_ to a node labelled from [*m*]_2_, when constructing the Bolthausen-Sznitman *n*-coalescent we will cut an edge on the path from the node labelled *i* to the root labelled 1 before the MRCA of [*m*] is reached, thus *i* would subtend the MRCA of [*m*]. The probability that a node labelled *i* ∈ [*n*]_*m*+1_ is not connected to a node labelled from [*m*]_2_ in 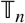 is

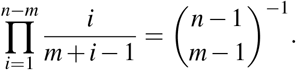

Even when there is no edge connecting a node labelled from [*n*]_*m*+1_ directly with a node labelled from [*m*]_2_, not all such paths of 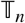 will contribute to 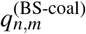. To contribute, we need that the cutting procedure does not lead to any *i* ∈ [*n*]_*m*+1_ being subtended by the MRCA of [*m*]. For the mentioned paths, this happens if and only if we cut all edges connecting nodes labelled from [*m*]_2_ to 1 before cutting any edge connecting 1 to nodes labelled from [*n*]_*m*+1_. We have Σ_*i*∈[*m*−1]_ *B_i_* edges adjacent to node 1, see the proof of Prop. 3. With the constraint that no edge connects a node labelled from [*n*]_*m*+1_ directly with a node labelled from [*m*]_2_, the sequential construction yields that, after relabelling, the nodes labelled with {1} ∪ [*n*]_*m*+1_ form a 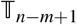 and thus there are 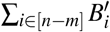 edges adjacent to the root of 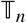 connecting to the nodes labelled with {1} ∪ [*n*]_*m*+1_, where 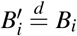 for independent 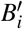. All edges adjacent to the node labelled 1 need to be cut before the MRCA of [*n*] is reached and they are cut at independent Exp(1) times. This means that the probability of cutting all edges connecting 1 to nodes labelled from [*m*]_2_ first is just drawing Σ_*i*∈[*m*−1]_ *B_i_* times without replacement from 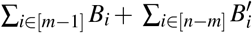 edges, where all Σ_*i*∈[*m*−1]_ *B_i_* edges connecting nodes labelled from to [*m*]_2_ have to be drawn. This probability equals

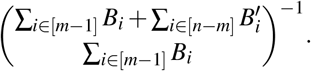

Integrating over all contributing paths of 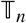 with the cutting constraint described above finishes the proof.

### 4.5 Proof of Thm. 1

#### Proof

We track the asymptotic frequencies (*P*_[*i*]_ (*t*))_*t*≥0_ of the ith biggest block for all *t* > 0 and 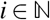. Consider a non-negative and measurable [0, ∞)-valued function *g* on the *k*-dimensional simplex

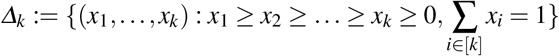

that is invariant under permutations (*x*_1_,…,*x*_*k*_) ↦ (*x*_*σ*(1)_,…,*x*_*σ*(*k*)_). [8, Thm. 1.2] shows that

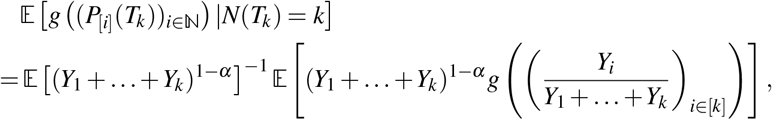

where *T_k_* is the waiting time until a state with ≤ *k* blocks is hit by the Beta-coalescent and *N*(*t*) is the number of blocks of *Π*_*t*_, thus we condition on the coalescent to hit a state with exactly *k* blocks.

We can apply this formula to compute 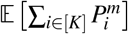 from Eq. (8), where *K* is the number of blocks at the last collision of the Beta-coalescent. For this, condition on *K* = *k*. With {*K*= *k*} = {*N*(*T*_*k*_) = *k*} ∩ {all blocks of *Π*_*T_k_*_ merge at the next merger}, the strong Markov property shows that the block frequencies at *T_k_* are independent of them merging at the next collision. However, these frequencies are, conditioned on *K*, just (*P_i_*)_*i*∈[*K*]_. For <u>*x*</u> ∈ *Δ_k_* we set 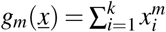 (which fulfills all necessary conditions to apply [8, Thm. 1.2]) and compute

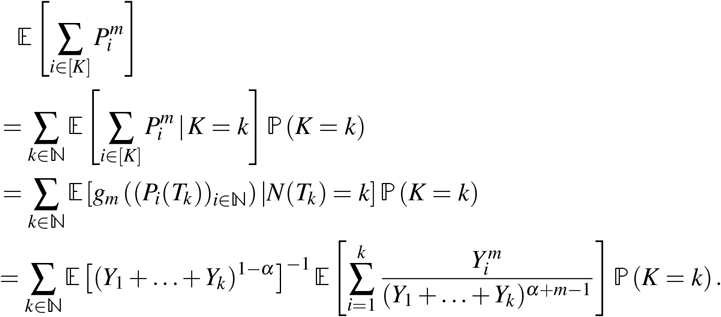

The distribution of *K* for the Beta-coalescent is known from [48, Thm. 3.5]. Using that

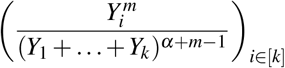

are identically distributed and 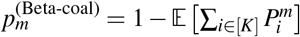 completes the proof.

### 4.6 Proof of Prop. 5

#### Proof

Consider any *Ξ*-coalescent (and its restrictions to [*n*], 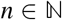). Since, for nested samples, 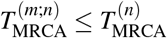 almost surely for any *n* ≥ *m*, we have

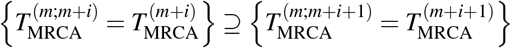

for any 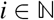. Thus, 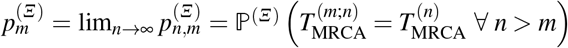 exists. Suppose first that the *Ξ*-coalescent comes down from infinity. Then, Eq. (8) shows 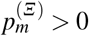. If the *Ξ*-coalescent stays infinite, *τ_m_* is almost surely finite, while 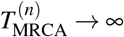 almost surely. Thus, 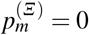.

Consider a *Ξ*-coalescent that neither comes down from infinity nor stays infinite. Then, 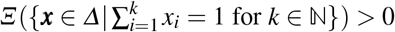. As stated in the introduction, in this case there is an almost surely finite waiting time *T* with *#Π_T_* < ∞ almost surely. Let *n_T_* be the finite number of blocks at time *T*. Again, exchangeability ensures, as in proving Eq. (7), that there is a positive probability that not all *i* ∈ [*m*] are in the same block of *Π_T_* (so in particular, with positive probability, 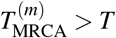). The strong Markov property of the *Ξ*-coalescent ensures that, given *n_T_*, *Π_T_* evolves like a *Ξ*-*n_T_*-coalescent, which can have at most *n_T_* mergers. In summary, with positive probability, more than one of the *n_T_* blocks at time *T* includes individuals from the subset [*m*] and the *n_T_* blocks are merged following a *Ξ*-*n_T_*-coalescent. Then, Eq. (7) shows that with positive probability, conditioned on the event that *k* > 1 blocks of *Π_T_* contain individuals from [*m*], also more than one block of the *Ξ*-coalescent at its last collision contains individuals of [*m*].

*Remark 2* Prop. 5 shows that 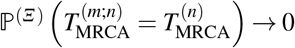 for fixed *m* and *n* → ∞ if the *Ξ*-coalescent stays infinite. The Bolthausen-Sznitman coalescent stays infinite [78, Example 15]; however, convergence to 0 is only of order 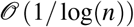.

### 4.7 Proof of recursions (4), (13), and (11)

The strong Markov property of a *Λ* -coalescent together with a natural coupling which we will introduce below allows us to describe many functionals of multiple-merger *n*-coalescents recursively by conditioning on their first jump, e.g. see [40] or [62]. We use this to prove recursions (4) and (11).

#### 4.7.1 Proof of Eq. (4)

Consider the probability 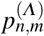 (see Eq. (4)) that a sample of size n shares the MRCA with a subsample of size *m* ∈ [*n* − 1]_2_. The boundary conditions *p*_*m,m*_ = 1 and *p*_*n*,1_ = 0 for *n* > 1 follow directly from the definition. We record how many individuals are merged at the first jump of the *n*-coalescent. Suppose a *k*-merger occurs which happens with probability *β*(*n*,*n*−*k*+1). Conditional on a *k*-merger, *ℓ* ≤ *m* of individuals that merge are taken from the subsample and *n* − *ℓ* are not with probability 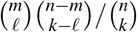, since the individuals that merge are picked uniformly at random without replacement. For 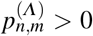, we need that not all m individuals are merged unless all n individuals are merged, thus *ℓ* < *m* or *k* = *n*. Writing *C*(*k*,*ℓ*) for the event that exactly *ℓ* lineages from the subsample are merged (with *ℓ* < *m* or *k* = *n*), the strong Markov property shows that

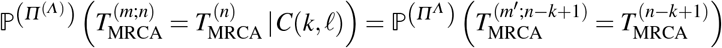

with 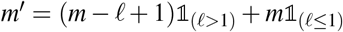, since among the ancestral lines (blocks) after the first collision, *m′* are subtended by the subsample. Summing over all possible values (recall the boundary conditions) yields recursion (4).

#### 4.7.2 Proof of Eq. (13)

We again condition on the event that *k* blocks are merged at the first jump. Only *k*-mergers where either all merged individuals are picked from the subsample [*m*] or none is sampled from [*m*] contribute positive probability to 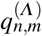. After the jump, we thus have *n* − *k* + 1 ancestral lineages present, from which either *m* − *k* + 1 or *m* are connected to the subsample. The strong Markov property and sampling without replacement for the *k*-merger then yields Eq. (13).

#### 4.7.3 Proof of Eq. (11)

Recall the natural coupling: if we restrict an n-coalescent with mutation rate *θ* to any *ℓ*-sized subset *L* ⊆ [*n*], the restriction is an *ℓ*-coalescent with mutation with the same rate *θ*. To prove recursion (11) we partition over three possible outcomes of the first event: it is a mutation on a lineage subtending the subsample (*E*_1_), it is a mutation on a lineage not subtending the subsample (*E*_2_), or it is a merger (*E*_3_). Naturally, before any mutation occurs, all edges are active.

We recall a few elementary facts. The time to the first mutation on any lineage is Exp(*θ*/2)-distributed (mutations on different/disjoint lineages are independent) and independent of the waiting time for the first merger. The minimum of independent exponential r.v.’s *X*_1_,…, *X*_*i*_ with parameters *α*_1_,…, *α*_*i*_ is again exponentially distributed with parameter 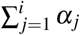. Finally, 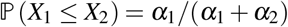.

The waiting times *X_i_* for events *E_i_* for 1 ≤ *i* ≤ 3 are all exponential; the one for *E*_1_ with rate *θm*/2, for *E*_2_ with rate *θ*(*n* − *m*)/2, and for *E*_3_ with rate *λ*(*n*). The probability of event *E*_1_ is 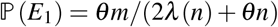 and, conditional on *E*_1_,

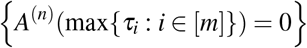

is determined by the *n* − 1 active lineages after the event. The memorylessness of the exponential distribution and natural coupling imply that after the first event, conditional on that event being *E*_1_, the remaining *n* − 1 lineages, of which (*m* − 1) subtend the subsample, follow an (*n* − 1)-coalescent with mutation rate *θ*. Thus,

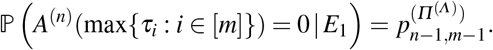

Analogously, we have 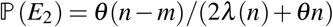. Given *E*_2_, we need to follow the coalescent of *n* − 1 lineages, of which *m* are from the subsample, which gives

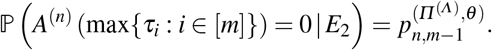

We have *P*(*E*3) = 1 − *P*(*E*_1_) − *P*(*E*_2_) = 2*λ*(*n*)/(2*λ*(*n*) + *θn*). To compute

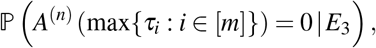

proceed exactly as in the proof of recursion (4) by partitioning over the number of mutant lineages involved in the merger, but with changed boundary conditions since 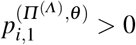, while 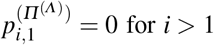 for *i* > 1. Summing over *E*_1_,*E*_2_,*E*_3_ yields Eq. (11).

### 4.8 Proof of Eq. (14)

Recall our assumption that block *π*_1_ always contains element 1. To see (14), we will show that, for *n* large enough,

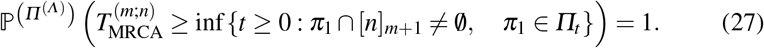

In words, the smallest block containing [*m*] appearing in the *n*-coalescent will always contain at least *m* + 1 elements; block [*m*] will almost never be observed. Hence, 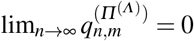.

Consider first *Λ* with 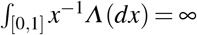, which makes the *Λ* -coalescent dust-free (no singleton blocks almost surely for *t* > 0) - see the proof of [70, Lemma 25]. For *t* > 0, [70, Prop. 30] shows that the partition block 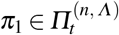 containing individual 1 at time *t* in the *Λ* -*n*-coalescent 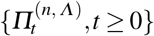 fulfills lim_*n*→∞_#*π*_1_/*n* > 0 almost surely. Thus, individual 1 has already merged before any time *t* > 0 if *n* > *N′*, where is *N′* a random variable on 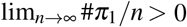 almost surely. However, within the subsample of fixed size *m*, we wait an exponential time with rate *λ* (*m*) for any merger of individuals in [*m*]. Thus, for n large enough individual 1 has almost surely already merged with individuals of [*n*]_*m*+1_ before merging with another individual in the subsample. Consider now *Λ* with 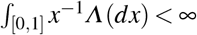, which shows that the coalescent has dust, i.e. there is a positive probability that there is a positive fraction of singleton blocks at any time *t*, see [70, Prop. 26]. In this case [37, Corollary 2.3] shows that at its first merger, for *n* → ∞, individual 1 merges with a positive fraction of all individuals 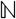 almost surely, which has to include individuals in [*n*]_*m*+1_. Since this is the earliest merger where the MRCA of [*m*] can be reached, the proof is complete.

Analogously we have the following:
**Corollary 1** *Consider any *Ξ*-coalescent which comes down from infinity and its restrictions to [*n*], 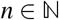. Fix subsample size 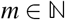. Let 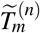 be the first time that any *i* ∈ [*m*] is involved in a merger in the *Ξ*-*n*-coalescentfor *n* ≥ *m*. We have*

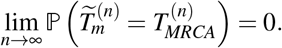

*Proof* If the *Ξ*-coalescent comes down from infinity, it fulfills 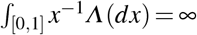, since it has to be dust-free. As above, we see that individual 1 has already merged before 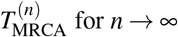 for *n* → ∞, which establishes the corollary.

## 5 Conclusion and open questions

By studying properties of nested samples we have aimed at understanding how much information about the evolutionary history of a population can be extracted from a sample, i.e. how the genealogical information increases if we enlarge the sample. In particular, we have focussed on multiple-merger coalescent (abbreviated MMC) processes derived from population models characterised by high fecundity and sweepstakes reproduction (abbreviated HFSR). In comparison with the Kingman-coalescent the general conclusion, at least for the statistics we consider, is that a subsample represents less well the ‘population’ or the complete sample from which the subsample was drawn when the underlying coalescent mechanism admits multiple mergers. The subsample reaches its most recent common ancestor (abbreviated MRCA; see Table 1 for definition of acronyms) sooner and shares less of the ancestral genetic variants (internal branches) with the complete sample under a MMC process than under the Kingman-coalescent. A similar conclusion can be broadly reached in comparison with exponential population growth. This seems to imply that one would need a larger sample for inference under a MMC than under a (time-changed) Kingman-coalescent. Large sample size has been shown to impact inference under the Wright-Fisher model [11], in particular if the sample size exceeds the effective size [86]. The main effect is that when sample size is large enough, one starts to notice multiple and/or simultaneous mergers in the trees ― events which would not be possible under the assumption that the sample size is fixed and the population size is arbitrarily large (and thus much larger than the sample size). The implication is that for any finite population, a large enough sample will ‘break down’ the coalescent approximation. One would also expect an impact of large sample size on inference under MMC.

The effective size in HFSR populations can be much smaller than in a Wright-Fisher population with the same census size [79,47,87]. Therefore, for almost any finite population, the genealogy of the whole population is not well approximated by the genealogy one derives under the assumption of a fixed sample size and an arbitrarily large population size. This therefore leaves the question of what one is making an inference about when one applies a coalescent-based inference method ― and how to evaluate whether the sample size is small enough that the coalescent approximation holds.

Naturally, this also means that our asymptotic results, most notably the representation for 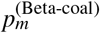 from Thm. 1, are not readily applied to real populations. Our asymptotic results rely on the assumption that the coalescent approximation holds for arbitrarily large sample sizes. However, our asymptotic results for 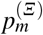 for any *Ξ* are still valid lower bounds for 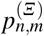 for any *n* until the coalescent approximation breaks down.

We focussed mostly on how genealogical properties (MRCA, internal branches) are shared between the complete sample and the nested subsample. These properties cannot be directly observed in genomic data, but they do reflect how much (and how old) polymorphisms can be potentially shared between the samples. We did not discuss similar quantities involving genetic variation (mutations) since, as we discuss in Sec. 2.2.1, a comparison of such quantities is confounded by the differences between different coalescent processes in the way time is measured.

All our results are applicable to a single non-recombining locus. A natural question to ask is if and how our results might change if we considered multiple unlinked loci. How would the statistics we consider, averaged over many unlinked loci, behave under multiple-merger coalescents in comparison with a (time-changed) Kingman-coalescent? DNA sequencing technology has advanced to the degree that sequencing whole genomes is now almost routine (see eg. [43, 4]). One could ask how large a sample from a HFSR population does one need to be confident to have sampled a significant fraction of the <u>genome-wide</u> ancestral variation? In this context, let 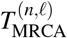 denote the TMRCA of the complete sample of size *n* at a non-recombining locus *ℓ* ∈ [*L*], and 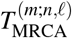 the TMRCA of a nested subsample of size *m* at same locus. Then we would like to compare the probability

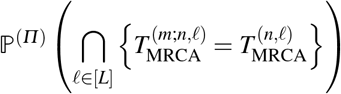

between different coalescent processes. And in fact, the independence of the genealogies at unlinked loci under the Kingman-coalescent, and Eq. (6), gives

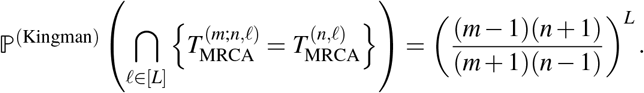

Under a multiple-merger coalescent process the genealogies at unlinked loci are not independent (see e.g. [32,15]).

We compared results from single-locus multiple-merger coalescent models with a time-changed Kingman-coalescent derived from a single-locus model of exponential population growth. Naturally one would like to compare results between genomic (multi-locus) models of HFSR with population growth to genomic models of HFSR without growth, and to genomic models of growth without HFSR. Some mathematical handle on the distributions of the quantities we simulated would (obviously) also be nice. However, these will have to remain important open tasks.

## Acknowledgements

We thank Alison Etheridge for many and very valuable comments and suggestions, especially regarding Theorem 1. BE was funded by DFG grant STE 325/17-1 to Wolfgang Stephan through Priority Programme SPP1819: Rapid Evolutionary Adaptation. FF was funded by DFG grant FR 3633/2-1 through Priority Program 1590: Probabilistic Structures in Evolution.

## A1 Population models

In this section we provide a brief overview of the population models behind the coalescent processes we consider, and why we think they are interesting. A detailed description of the coalescent processes is given in Sec. A2.

A universal mechanism among all biological populations is reproduction and inheritance. Reproduction refers to the generation of offspring, and inheritance refers to the transmission of information necessary for viability and reproduction. Mendel’s laws on independent segregation of chromosomes into gametes describe the transmission of information from a parent to an offspring in a diploid population. For our purposes, however, it suffices to think of haploid populations where one can think of an individual as a single gene copy. By tracing gene copies as they are passed on from one generation to the next one automatically stores two sets of information. On the one hand one stores how frequencies of genetic types change going forwards in time; on the other hand one keeps track of the <u>ancestral</u>, or genealogical, relations among the different copies. This <u>duality</u> has been successfully exploited for example in modeling selection [34,35]. To model genetic variation in natural populations one requires a mathematically tractable model of how genetic information is passed from parents to offspring. In the Wright-Fisher model offspring choose their parents independently and uniformly at random. Suppose we are tracing the ancestry of *n* ≥ 2 gene copies in a haploid Wright-Fisher population of *N* gene copies in total. For any pair, the chance that they have a common ancestor in the previous generation is 1/*N*. Informally, we trace the <u>genealogy</u> of our gene copies on the order of 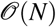 generations until we see the first <u>merger</u>, i.e. when at least 2 gene copies (or their ancestral lines) find a <u>common ancestor</u>.

If *n* is small relative to *N*, when a merger occurs, with probability 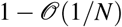 it involves just two ancestral lineages. This means that if we measure time in units of *N* generations, and assume *N* is very large, the random ancestral relations of our sampled gene copies can be described by a continuous-time Markov chain in which each pair of ancestral lines merges at rate 1 and no other mergers are possible. We have, in an informal way, arrived at the Kingman-coalescent [56,58,57]. One can derive the Kingman-coalescent not just from the Wright-Fisher model but from any population model which satisfies certain assumptions on the offspring distribution [61,71,64]. These assumptions mainly dictate that higher moments of the offspring number distribution are small relative to (an appropriate power of) the population size. The Kingman-coalescent, and its various extensions, are used almost universally as the ‘null model’ for a gene genealogy in population genetics. The Kingman-coalescent is a remarkably good model for populations characterised by low fecundity, i.e. whose individuals have small numbers of offspring relative to the population size.

The classical Kingman-coalescent is derived from a population model in which the population size is constant between generations. Extensions to stochastically varying population size, in which the population size does not vary ‘too much’ between generations, have been made [54]; the result is a time-changed Kingman-coalescent. Probably the most commonly applied model of deterministically changing population size is the model of exponential population growth (see eg. [25,41,30]). In each generation the population size is multiplied by a factor (1 + *β/N*), where *β* > 0. Therefore, the population size in generation k going forward in time is given by *N_k_* = *N*(1 + *β*/*N*)^*k*^ where *N* is taken as the ‘initial’ population size. It follows that the population size ⌊*Nt*⌋ generations ago is *Ne^−βt^*. [30] show that exponential population growth can be distinguished from multiple-merger coalescents (in which at least three ancestral lineages can merge simultaneously), derived from population models of high fecundity and sweepstakes reproduction, using population genetic data from a single locus, provided that sample size and number of mutations (segregating sites) are not too small.

A diverse group of natural populations, including some marine organisms [46], fungi [1,80,51], and viruses [84] are highly fecund. By way of example, individual Atlantic codfish [60,68] and Pacific oysters [59] can lay millions of eggs. This high fecundity counteracts the high mortality rate among the larvae (juveniles) of these populations (Type III survivorship). The term ‘sweepstakes reproduction’ has been proposed to describe the reproduction mode of highly fecund populations with Type III survivorship [45]. Population models which admit high fecundity and sweepstakes reproduction (HFSR) through skewed or heavy-tailed offspring number distributions have been developed [64,65,79,31,73,53]. In the haploid model of [79], each individual independently contributes a random number *X* of juveniles where (*C*, *α* > 0)

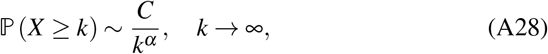

and *x_n_* ~ *y_n_* means *x_n_/y_n_* → 1 as *n* → ∞. The constant *C* > 0 is a normalising constant, and the constant α determines the skewness of the distribution. The next generation of individuals is then formed by sampling (uniformly without replacement) from the pool of juveniles. In the case α < 2 the random ancestral relations of gene copies can be described by specific forms of <u>multiple-merger</u> coalescent processes [72]. We remark that the fate of the juveniles need not be correlated to generate multiple-mergers in the genealogies ― the heavy-tailed distribution of juveniles means that occasionally one ‘lucky’ individual contributes a huge number of juveniles while all others contribute only a small number of juveniles. Uniform sampling without replacement from the pool of juveniles means that the lucky individual leaves significantly more descendents in the next generation than anyone else, and this is what generates multiple mergers of ancestral lines.

Coalescent processes derived from population models of HFSR (see (A28) for an example) admit multiple mergers of ancestral lineages [24,70,71,76,65,72,63]. Mathematically, we consider exchangeable n-coalescent processes, which are Markovian processes 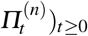 on the set of partitions of [*n*]: = {1, 2,…, *n*} whose transitions are mergers of partition blocks (a ‘block’ is a subset of [*n*], see Sec. A2) with rates specified in Sec. A2. The blocks of 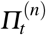 show which individuals in [*n*] share a common ancestor at time t measured from the time of sampling. Thus, the blocks of 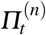 can be interpreted as ancestral lineages. The specific structure of the transition rates allows to treat a multiple-merger *n*-coalescent as the restriction of an exchangeable Markovian process (*Π_t_*)_*t*_≥_0_ on the set of partitions of 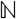, which is called a multiple-merger coalescent (abbreviated MMC) process. MMC processes are referred to as *Λ*-coalescents (*Λ* a finite measure on [0,1]) [24,70,71] if any number of ancestral lineages can merge at any given time, but only one such merger occurs at a time. By way of an example, if 1 ≤ *α* ≤ 2 in (A28) one obtains a so-called Beta(2 − *α, α*)-coalescent [72] (Beta-coalescent, see Eq. (A35)). Processes which admit at least two (multiple) mergers at a time are referred to as *Ξ* -coalescents (*Ξ* a finite measure on the infinite simplex *Δ*) [76,64,65]. See Sec. A2 for details. Specific examples of these MMC processes have been shown to give a better fit to genetic data sampled from Atlantic cod [12,18,2,16,19] and Japanese sardines [67] than the classical Kingman-coalescent. See e.g. [29] for an overview of inference methods for MMC processes. [46] review the evidence for sweepstakes reproduction among marine populations and conclude ‘that it plays a major role in shaping marine biodiversity’.

MMC models also arise in contexts other than high fecundity. [17] show that repeated strong bottlenecks in a Wright-Fisher population lead to time-changed Kingman-coalescents which look like *Ξ* -coalescents. [27,28] show that the genealogy of a locus subjected to repeated beneficial mutations is well approximated by a *Ξ* -coalescent. [75] provides rigorous justification of the claims of [66,22] that the genealogy of a population subject to repeated beneficial mutations can be described by the Beta-coalescent with *α* = 1 (also referred to as the Bolthausen-Sznitman coalescent [20]). These examples show that MMC processes are relevant for biology. We refer the interested reader to e.g. [10,25,5,33,9,13] for a more detailed background on coalescent theory.

## A2 Coalescent processes

To keep our presentation self-contained a precise definition of the coalescent processes we will need will now be given. We follow the description of [19]. A coalescent process *Π* is a continuous-time Markov chain on the partitions of 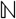. Let *Π*^(*n*)^ denote the restriction to [*n*], and write 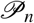 for the space of partitions of [*n*]. A partition 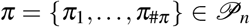 has #*π* blocks which are disjoint subsets of [*n*]. We assume the blocks *π_i_* are ordered by their smallest element; therefore we always have 1 ∈ *π*_1_. In general a merging event can involve r distinct groups of blocks merging simultaneously. We write <u>*k*</u> = (*k*_1_,…, *k_r_*) where *k*_i_ ≥ 2 denotes the number of blocks merging in group *i*. Here *r* ∈ [⌊*#π*/2⌋], *k*_1_ +⋯+*k_r_* ∈ [*#π*]_2_ and 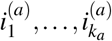 will denote the indices of the blocks in the ath group. By *π′* ≺_*#π*,<u>*k*</u>_ *π* we denote a transition from *π* to *π′* = *A* ∪ *B* where

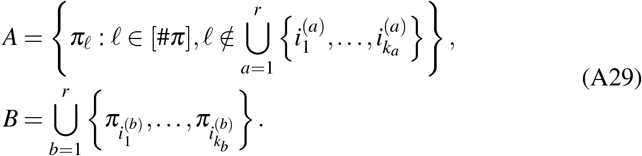

In (A29), set A (possibly empty) contains the blocks not involved in a merger, and *B* lists the blocks involved in each of the r mergers. By *π′ ≺#_π,k_ π* we denote the transition in a *Λ* -coalescent where *k* ∈ [*#π*]_2_ merge in a single merger and *π′* is given as in (A29) with *r* = 1; ie. only one group of blocks merges in each transition. By *π′ ≺_#π_ π* we denote a transition in the Kingman-coalescent where *r* = 1 and 2 blocks merge in each transition.

Now that we have specified the possible transitions, we can state the rates of the transitions. Let Δ denote the infinite simplex *Δ* = {(*x*_1_, *x*_2_,…): *x*_1_ ≥ *x*_2_ ≥ … ≥ 0,Σ_i_x_i_ ≤ 1}; let ***x*** denote an element of *Δ*. Define the functions *f* (***x***;*#π,<u>k</u>*) and *g*(***x**;#π,<u>k</u>*) on *Δ*_**0**_: = *Δ* \ {(0, 0,…)} where 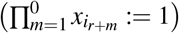, and *s* = *#π* − *k*_1_ − … − *k_r_*, by

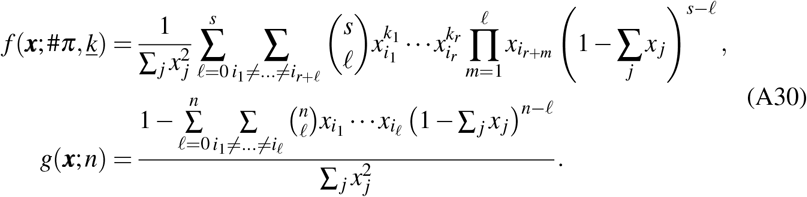

where *x_i_0__*: = 1. For a finite measure *Ξ* on *Δ*, set *Ξ*_0_: = *Ξ*(·∩*Δ*_0_) and *a*: = *Ξ*({(0, 0,…)}).

Then, define

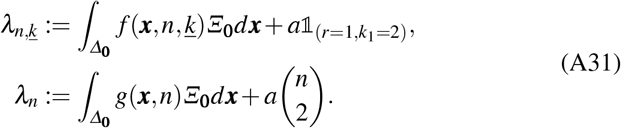

A *Ξ*-coalescent [76] is a continuous-time 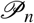-valued Markov chain with transitions *q_π, π′_* given by, where *λ_n,<u>k</u>_* and *λ_n_* are given in (A31),

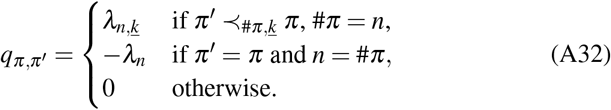

A *Λ*-coalescent [24,70,71] is a specific case of a *Ξ*-coalescent where *Ξ**0*** only has support on 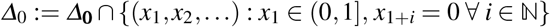 [76]. Let *Λ* denote the restriction of *Ξ* on its first coordinate (which makes *Λ* a finite measure on [0, 1]). The transition rate of *π′ ≺_π,k_ π* becomes, where *#π* = *n*, 2 ≤ *k* ≤ *n*,

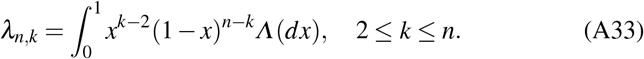

The total rate of *k*-mergers in a *Λ* -coalescent is given by 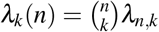 for 2 ≤ *k* ≤ *n*. The total rate of mergers given *n* ≥ 2 active blocks is

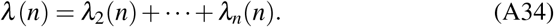

An important example of a *Λ* -coalescent is the Beta(2 − *α*, *α*)-coalescent [79] where the *Λ* measure is associated with the beta density, where ***B***(·, ·) is the beta function,

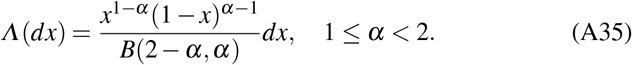

The total rate of a *k*-merger 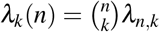 (see Eq. (A33)) is then given by, for 2 ≤ *k* ≤ *n*,

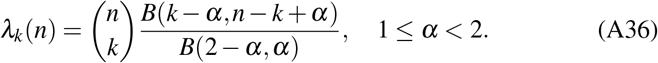

For *α* = 1 the Beta(2 − *α*, *α*)-coalescent is the Bolthausen-Sznitman coalescent [20,39]. The Beta-coalescent is well-studied, there are connections to superprocesses, continuous-state branching processes (CSBP) and continuous stable random trees as described e.g. in [14] and [7].

## A3 Goldschmidt and Martin’s construction of the Bolthausen-Sznitman *n*-coalescent

From [39], we recall the construction of the Bolthausen-Sznitman n-coalescent by cutting the edges of a random recursive tree. Let 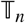 be a random recursive tree with *n* nodes. We can construct 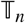 sequentially as follows

i. Start with a node labelled with 1 (the root) and no edges,
ii. If *i* < *n* nodes are present, add a node labelled with *i* + 1 and one edge connecting it to a node in [*i*] picked uniformly,
iii. stop if *n* nodes are present.

The object 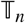 is a labelled tree, each node has a single label. We consider a realisation of 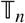 and transform this tree over time into labelled trees with fewer nodes with nodes amassing multiple labels.
i. Each edge of 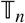 is linked to an exponential clock. The clocks are i.i.d. Exp(1)-distributed.
ii. We wait for the first clock to ring. At this time, we cut/remove the edge whose clock rang first. The tree is thus split in two trees, one of these trees includes the node with label 1. We denote this tree by 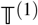, the other tree by 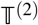. Let *e*_1_ be the node of 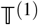 that was connected to the removed edge.
iii. All labels of 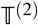 are added to the set of labels of *e*_1_. Remove 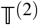 including its clocks.
iv. Repeat from (ii), using 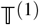 labelled as in (iii) with the (remaining) clocks from (i). Stop when 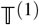 in step (iii) consists of only a single node and no edges.
v. For any time *t*, label sets at the nodes of 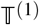 (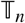 before the first clock has rang) give a partition 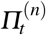 of [*n*]. The process 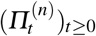 is a Bolthausen-Sznitman *n*-coalescent (set 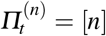 if *t* is bigger than the time at which we stopped the cutting procedure).

**Fig. 7.**
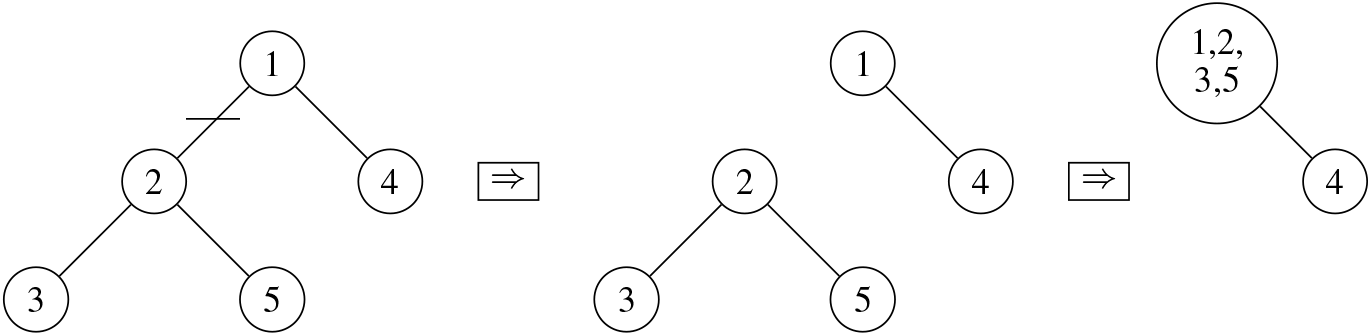
Example for the first cutting and relabelling step (ii), (iii) for the construction from [39].

Figure A3 shows an illustration of steps (i)-(iii) for a realisation of 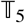.

